# Fast and slow synaptic plasticity enables concurrent control and learning

**DOI:** 10.1101/2024.09.06.611710

**Authors:** Brendan A. Bicknell, Peter E. Latham

## Abstract

During many tasks the brain receives real-time feedback about performance. What should it do with that information, at the synaptic level, so that tasks can be performed as well as possible? The conventional answer is that it should learn by incrementally adjusting synaptic strengths. We show, however, that learning on its own is severely suboptimal. To maximize performance, synaptic plasticity should also operate on a much faster timescale – essentially, the synaptic weights should act as a control signal. We propose a normative plasticity rule that embodies this principle. In this, fast synaptic weight changes greedily suppress downstream errors, while slow synaptic weight changes implement statistically optimal learning. This enables near-perfect task performance immediately, efficient task execution on longer timescales, and confers robustness to noise and other perturbations. Applied in a cerebellar microcircuit model, the theory explains longstanding experimental observations and makes novel testable predictions.

## Introduction

The standard view of learning is that synaptic strengths are adjusted to minimize some loss – typically the performance on a task, or set of tasks (Richards and Kording 2023; Bredenberg and Savin 2023). From this perspective, given the complexity of the nervous system and the range of tasks it must perform, conventional wisdom is that synaptic strengths change slowly. This makes sense for tasks with a discrete set of responses: Do I lick left or right in response to a stimulus? Is that grating oriented to the right or left of vertical? However, for naturalistic tasks with continuous output, such as reaching for an object, feedback about performance could be used to adjust weights on much faster timescales. In fact, for sufficiently fast feedback compared to the timescale over which the world changes, fast weight changes could, at least in principle, enable near-perfect performance.

Here we propose that synapses can maximize task performance by adjusting weights on two timescales. On fast timescales, synapses can use real-time feedback to suppress immediate errors. As long as the world changes relatively smoothly over time, if feedback at one moment indicates that neural output is too high or low, synaptic strengths can be transiently decreased or increased to compensate. On slow timescales, synapses can use the same feedback signal for statistically optimal learning.

We first illustrate the main concept with a toy model, demonstrating how two timescales of plasticity can work in concert. We then consider a more realistic model of a neuron, where synapses dynamically integrate input and feedback to extract useful learning signals from noisy observations. We derive synaptic update rules for this model by framing synaptic plasticity as an optimal control problem. This leads to substantial improvements over classical gradient-based learning. We then generalize the theory to incorporate small populations of neurons and delayed feedback transmitted via spikes. Applied to a model of temporal processing in the cerebellum, our theory provides normative explanations for common experimental observations, and makes novel testable predictions. Altogether, these results provide a principled account of how the brain can exploit multiple timescales of plasticity for efficient online adaptation and learning.

## Results

We propose that neurons can optimize their performance by having two separate timescales of synaptic plasticity. To illustrate the general idea, consider a simple online linear regression problem. We model a single neuron that must learn to transform a set of time-varying input rates, *ν*_*i*_(*t*), into a target output, *y*\*(*t*). The output of the neuron is given by

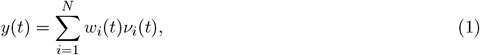

with parameters *w*_*i*_ denoting its tunable synaptic weights. The classical solution to this problem is to discretize time and iteratively modify the weights using the delta rule (Widrow and Hoff 1960),

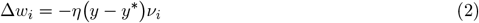

where Δ*w*_*i*_ ≡ *w*_*i*_(*t* + Δ*t*) − *w*_*i*_(*t*). This eventually minimizes the output error provided the learning rate, *η*, is small.

The delta rule is attractive because it is generally effective and it is biologically plausible – each synapse requires access only to its own local input rate, *ν*_*i*_, and a global error feedback signal, *y* − *y*\*. If, however, *ν*_*i*_ and target output, *y*\*, change smoothly over time, temporal correlations can be exploited to dramatically improve performance. To exploit temporal correlations, we endow the synaptic weights with independent fast and slow components,

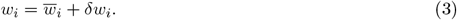

The role of the fast weights, *δw*_*i*_, is to transiently adjust neuronal output to suppress immediate errors – errors that can be predicted if *y*\* changes slowly. The role of the slow weights, *w*_*i*_, is to adapt, on a much slower timescale, to the true weights, thereby reducing the need for ongoing fast-weight corrections (Fig. 1a).

**Figure 1:**
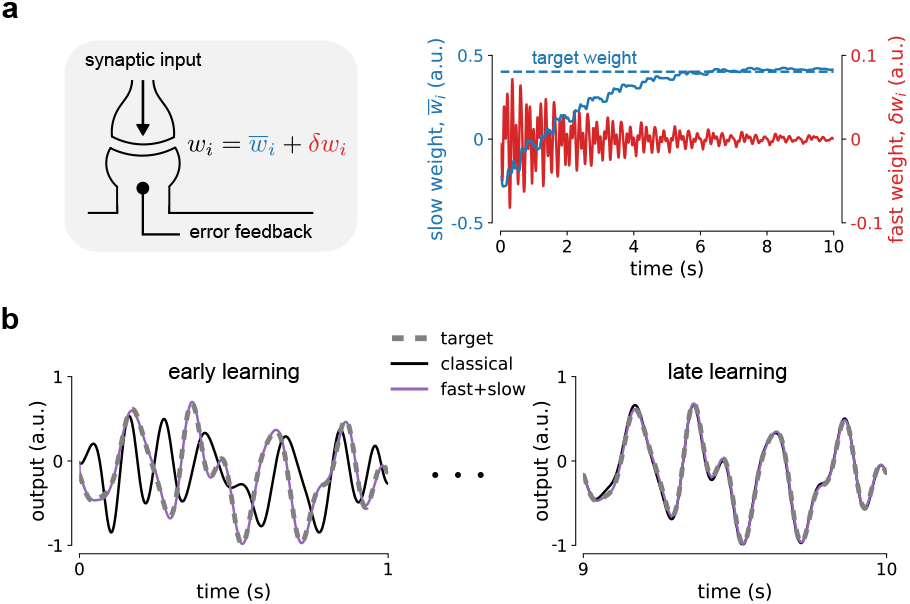
A theory of fast and slow synaptic plasticity. **a)** Left: A synapse must make online adjustments to its strength by integrating local signals, such as its own input and error feedback. We propose that these signals can be optimally exploited through two timescales of plasticity. ‘Fast weights’, *δw*_*i*_, fluctuate rapidly to suppress downstream error, whereas ‘slow weights’, *w*_*i*_, converge gradually to the values required of a given task. Right: In the toy-model simulation from panel (b), as the slow weights find the solution, fast-weight fluctuations are reduced. Shown are example weight trajectories from one randomly selected synapse out of 20. **b)** In an illustrative online regression task, a neuron must learn to match its output to a time-varying target (gray dashed line). With a classical delta rule (black line), weights adapt over time to eventually correct the output. With two timescales of plasticity (purple line), fast weights can pin the output to the target from the outset, while slow weights evolve in the background to learn the task.

For the model described by equation (1), a simple implementation of this strategy is to observe the error at discrete times *t*, and then update all of the fast weights uniformly via

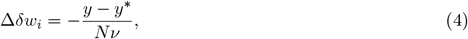

where Δ*δw*_*i*_ ≡ *δw*_*i*_(*t* + Δ*t*) − *δw*_*i*_(*t*) and 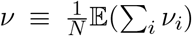 is the expected input rate per synapse (computed via a time average, or assumed as a known parameter). Summed over synapses, the fast weights subtract the observed error from the output, and straightforward algebra gives us

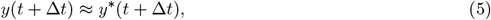

where the approximation is good if there are a large number of synapses and Δ*t* is small relative to how fast the target and input firing rates change (*Methods: Online linear regression*).

While suppressing error with rapidly fluctuating weights is effective, continually making these shortterm adjustments is likely to be energetically costly. This can be addressed by allowing the slow weights to learn. Intuitively, the closer the output is to the target, the less work will be needed to correct it. Seeking to make long-term adjustments to the slow weights, such that, ultimately, 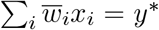, leads to local updates,

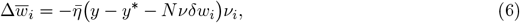

where 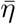 is a small learning rate (*Methods: Online linear regression*). This is similar to the usual delta rule, but with the fast weights subtracted off: it’s not hard to see that 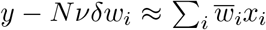. Slow-weight learning is thus driven by a modified error signal that represents what would have been observed in the absence of all of the fast corrections, isolating the contribution of 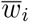. This coupling between the two update rules makes them compatible: so long as the slow weights know what the fast weights are doing, greedily suppressing error does not come at the expense of learning – even though the error is a crucial teaching signal.

In simulations of this simple strategy, the weights that solve the regression problem can be learned just as efficiently as in the classical case, but with the striking difference that the output is pinned to the target from the outset (Fig. 1b). Thus, a neuron, or indeed an animal, could accurately execute a task even while it is still being learned. Below we develop a control theory framework that generalizes this to more realistic scenarios.

### Synaptic plasticity as optimal control

We now apply this idea to a more realistic setting: a model neuron driven by multiple spike trains and subject to intrinsic noise. Similar to the regression problem above, the neuron must tune its synaptic weights to drive a downstream output, *y*, to match a time-varying target, *y*\*. The target could represent, for instance, the movement of a limb, but in this analysis we will take it to be an abstract, time-varying one-dimensional signal. We assume there are *N* synapses, and each one has access to two sources of information: the presynaptic spikes it receives, and an error signal.

The dynamics of the model neuron are described by equations for the total synaptic current, *I*, and firing rate, *r*,

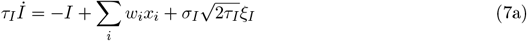

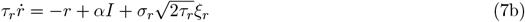

where a dot denotes a time derivative. In these equations, *τ*_*I*_ ∼ 5 ms and *τ*_*r*_ ∼ 50 ms denote the synaptic and rate-modulation time constants, *w*_*i*_ denotes the weight of synapse *i*, and each *x*_*i*_ is an independent Poisson spike train represented by a sum of delta functions,

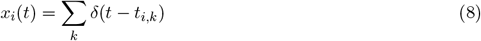

where *t*_*i,k*_ is the time of the *k*^th^ spike arriving at synapse *i*. The parameter *α* sets the scale of firing rate response to synaptic current, and the final terms, *ξ*_*I*_ and *ξ*_*r*_, denote zero-mean, unit-variance Gaussian white noise processes: ⟨*ξ*_*I*_(*t*)*ξ*_*I*_(*t*′)⟩ = *δ*(*t* − *t*′), and similarly for *ξ*_*r*_. Functionally, a barrage of input spikes is linearly filtered to drive the firing rate of the neuron.

The downstream output of the neuron, *y*, is driven by the firing rate of the neuron and white noise,

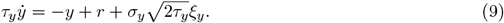

The time constant, *τ*_*y*_ (taken to be ∼100 ms), determines how quickly the output responds to changes in firing rate, and the final term is, as above, zero-mean, unit-variance Gaussian white noise.

We develop the theory using a teacher-student learning framework: the target, *y*\*, is generated by simulating equations (7) and (9), except with no noise and with a set of target weights, denoted 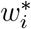, in place of *w*_*i*_. The target weights drift slowly around a mean value, *µ*_*w*_, with time constant *τ*_*w*_ ∼ 10^3^ s, modeling environmental variability or ongoing changes in the rest of the brain (Aitchison et al. 2021). We assume, for simplicity, that an instantaneous, continuous error signal is provided to all synapses, such as by diffuse transmission of a neuromodulator (Magee and Grienberger 2020); below we will relax this assumption, and incorporate delayed feedback communicated by spikes. The error signal is modeled as

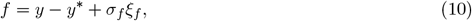

where *ξ*_*f*_ is zero-mean, unit-variance Gaussian white noise.

The full derivation of the plasticity rule is provided in *Methods: Derivation of the Bayesian plasticity rule*. Here we outline the general approach and main results. Similar to the analysis above, we decompose the synaptic weights into independent fast and slow components, 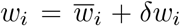, as in equation (3). Framing synaptic plasticity as an optimal control problem, we find that by deriving fast-weight dynamics that minimize the error we simultaneously obtain a highly effective slow-weight learning rule.

To derive the fast-weight rule, we consider a typical control strategy: using past error signals (equation (10)), compute a control variable, 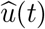, such that applying it as an input to the system (in our case, at the level of synaptic currents) will minimize future output error. Analogous to the toy model above, rather than invoke an external control system for this purpose, we assume everything is implemented locally at individual synapses.

Suppose that each synapse can compute such an error-minimizing control, 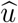. Setting the fast weights uniformly as

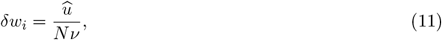

with *ν* denoting the expected input rate per synapse, then leads to 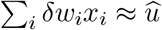.

We would like to choose 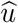 to minimize the squared error, (*y* − *y*\*)^2^. However, there is a cost to making the squared error very small: 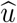, and thus *δw*_*i*_, will undergo large fluctuations. To navigate the tradeoff between minimizing the squared error while making sure the fast-weight fluctuations aren’t so large, we find 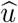 via

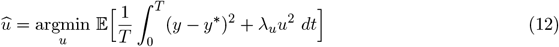

where *T* is the task duration and the expectation, 𝔼 […], is over the noise. The parameter *λ*_*u*_ is the ‘cost of control’; making it larger reduces fluctuations in the fast weights but increases the squared error between *y* and *y*\*; making it smaller has the opposite effect. The quadratic loss in equation (12), along with the Gaussian white noise in the dynamical equations, means that 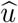 can be computed as a solution to the classic linear-quadratic-Gaussian (LQG) control problem. This approach results in a dynamical system that performs a Bayes-optimal estimate of unobserved states (such as the true underlying error, *y* −*y*\*), and uses that to construct an optimal linear controller (Crassidis and Junkins 2011).

The resulting equations predict how synapses should process input and feedback for optimal plasticity. For ease of intuition, we present the results here with some simplifying approximations. The results without approximations, given by equation (76), are used in simulations, although the approximate equations below provide comparable performance (Fig. S1).

To set the fast weights, *u* is computed by processing the error feedback signal, *f* (equation (10)), via

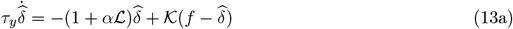

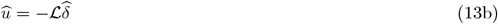

(*Methods*, equations (80a) and (92)). Equation (13a) filters the noisy feedback to compute an online estimate, 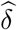, of the true error, *δ* = *y* − *y*\*, using an optimal feedback gain, 𝒦. The control variable is designed to negate the estimated error, with the magnitude of corrections determined by the optimal control gain, ℒ (equation (13b)). The gains 𝒦 and ℒ depend on the model parameters, reflecting both the integrative dynamics of the neuron and downstream output, and the statistics of the noise.

How do the fast weights evolve in this scenario? Taking a time derivative of equation (11), and substituting equations (13a) and (13b), leads to

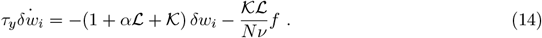

The fast weights are thus a low-pass filtered version of the negative of the error signal, which is the optimal strategy in the presence of noise. To see how this relates to the toy-model strategy above, we use the fact that when the control cost, *λ*_*u*_, and feedback noise variance, 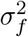, are small, the optimal gains ℒ and 𝒦 are large (*Methods*, equations (87) and (91)). In this regime, the last term on the right-hand side of equation (14) dominates the dynamics; if we consider discrete updates with a time step Δ*t*, equation (14) becomes

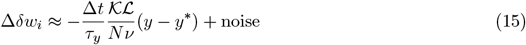

where we used equation (10) for *f*. Choosing the control cost to ensure that the product 𝒦ℒ obeys the relationship 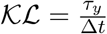 and ignoring the noise, we recover the fast-weight updates for the toy model (equation (4)). Details aside, the key principle is the same: observe the current error, then greedily adjust weights to compensate.

What about the slow weights – should they even be updated? And if so, how? To answer the first question, we note that much of the error being controlled by the fast weights is due to mismatch between the slow weights, 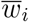, and the target weights, 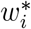. Which makes sense: in the absence of noise, when 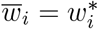 the weights do not need to be adjusted. That can be seen for the toy model in the right panel of Fig. 1a; for the more realistic model it’s shown explicitly in *Methods*, equation (48a). This tells us that the closer the slow weights are to their target values, the smaller the required control signal. Thus, given there is always a maximum control that can be applied, pushing the slow weights towards the target weights will improve performance.

A simple approach to updating the slow weights would be to use a modified delta rule, similar to equation (6). However, in this more realistic setting, the delta rule is suboptimal for two reasons. First, the error feedback signal, *f*, is noisy, which leads to noisy weight fluctuations, and even instability unless the learning rate is very small. Second, as pointed out by Aitchison et al. (2021), efficient weight updates should depend on uncertainty, with more uncertain weights updated more rapidly.

Qualitatively at least, the first problem can be fixed by filtering the feedback signal, *f*, and the second by scaling the learning rate by the uncertainty. Determining exactly how to design the filter and how much to scale, is, however, somewhat nontrivial. Fortunately, the analysis used to derive the update rule for the fast weights leads naturally to a rule for updating the slow weights that addresses these problems.

As shown in *Methods: Derivation of the Bayesian learning rule* (see in particular equation (80)), the update rule for the slow weights is

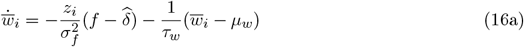

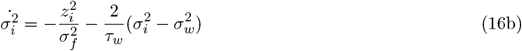

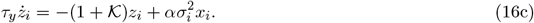

The first term on the right-hand side of equation (16a) looks very similar to the delta rule. However, there are three notable differences. First, the term proportional to 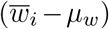 leads to relaxation to the target-weight mean in the absence of feedback. This reflects the fact that, as mentioned above, the weights drift on a timescale of *τ*_*w*_, so without feedback the mean, *µ*_*w*_, becomes the optimal estimate of the weight. Second, the raw error feedback signal, *f*, is replaced by a prediction error, 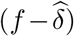. Because 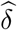 is just a filtered version of *f* (equation (13a)), 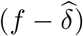 reflects deviations of the error signal about its time-averaged value. Third, and most important, the effective learning rate is proportional to *z*_*i*_, which plays the role of an eligibility trace, so is different for every synapse. Equation (16c) tells us that *z*_*i*_ increases with the variable 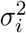, which reflects weight uncertainty, so the higher the uncertainty the higher the learning rate – as expected. Determining the dependence of 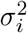 on the firing rate is not completely straightforward, but we show in *Methods* (see in particular equation (99)) that it scales as 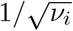 where, recall, *ν*_*i*_ is the rate of input to synapse *i*. This too is expected, at least qualitatively: the higher the firing rate, the more certain a synapse is about its value, and the lower the learning rate.

Mathematically, equations (16a) and (16b) represent the evolving mean and variance of a posterior distribution over target weights, similar to the approach introduced by Aitchison et al. (2021). This is the best estimate that can be formed by the synapse, given the observed data, and therefore the most efficient learning rule for 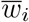. To highlight this qualitative difference from classical counterparts, and make connections to previous work, we will refer to our plasticity rule and its generalizations as ‘Bayesian plasticity’, in line with Aitchison et al. (2021).

Using the teacher-student task, we compared our plasticity rule to a classical gradient-based rule for a neuron with 1000 synapses. The results are shown in Fig. 2. The classical approach is a generalized delta rule based on the Real-Time Recurrent Learning algorithm (Williams and Zipser 1989; Pearlmutter 1995), which is the canonical approach for online learning in dynamic models. We included simulations of the full Bayesian rule, as well as versions using the slow and fast components alone. Learning with the Bayesian slow-weight rule alone is already superior to the classical rule, both in terms of total output error (Fig. 2a) and the speed of convergence of the weights towards their target values (Fig. 2b). Output error is reduced further still when the fast weights are included, due to suppression of noise and compensation for weight mismatch at early stages of learning. Moreover, weight convergence in this case is almost indistinguishable from when slow-weight learning is implemented in isolation. Although a similarly low level of error is obtained using the fast weights alone, this requires much larger control currents (Fig. 2c), and no memory of the task is actually retained in the weights. With some small modifications (*Methods: Feedback delays*), our rule also accounts for time lags in the feedback signal, representing communication delays in the brain. In this case, slow-weight learning is virtually unaffected and fast-weight control remains effective for delays on the order of the downstream time constant (Fig. 2d).

**Figure 2:**
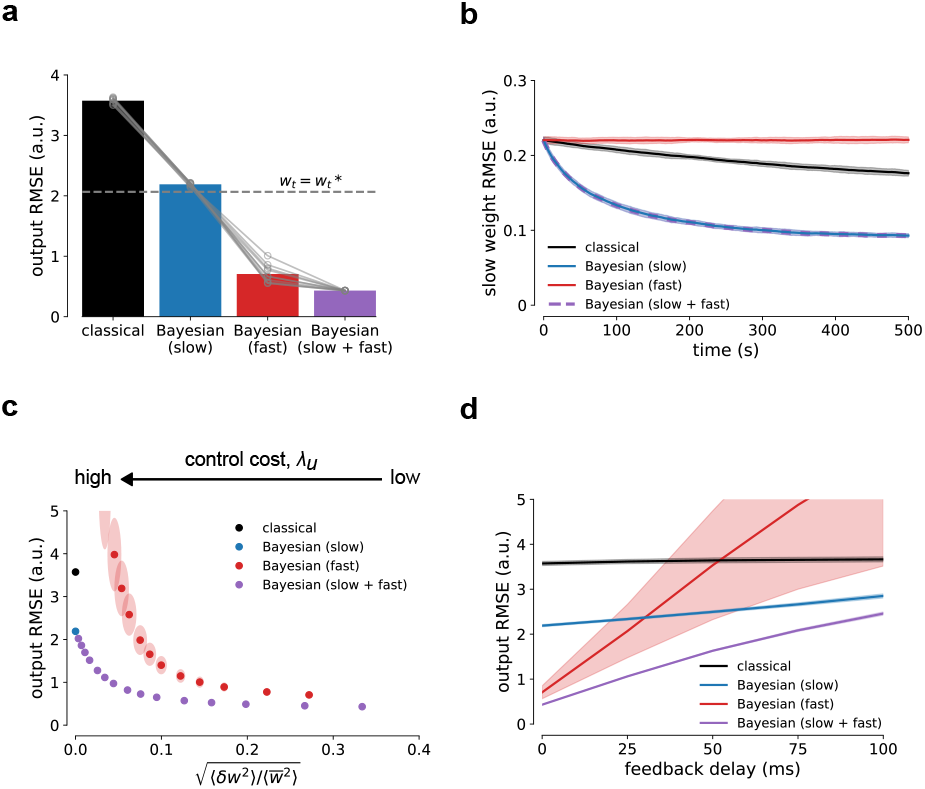
Synaptic plasticity as optimal control. **a)** Performance comparison between the classical gradient-based rule (black) and Bayesian rule without fast weights (blue), with plastic fast weights and frozen slow weights (red), and with plastic slow and fast weights (purple). Bars denote root-mean-squared error (RMSE) between output, *y*, and target output, *y*\*, averaged over the entire task. Data points denote different random seeds. The dashed line gives the average error when slow weights are set to their target values at every point in time. The learning rate for the classical rule and control cost for the Bayesian rule were selected via grid search to minimize output error. **b)** The Bayesian rules yield faster convergence of weights to their target values compared to the classical rule, quantified as RMSE between weight vectors. Shaded areas are standard deviation from 10 random seeds. **c)** Root-mean-squared output error versus fast weight fluctuations, the latter computed as the root-mean-squared fast weight divided by root-mean-squared slow weight (averages taken over entire task). The size of the fast weight fluctuations were controlled by the cost parameter *λ*_*u*_ (see equation (12)). In the full plasticity rule (purple), even small fluctuations of ∼ 10% lead to a large reduction in error. Without slow-weight learning (red), much more control is needed for a similar level of performance. Points denote the average over seeds for a fixed *λ*_*u*_; the shaded ellipses denote standard deviation across seeds. **d)** Control is effective for feedback delays shorter than the output time constant (here, *τ*_*y*_ = 100 ms). Shaded areas are standard deviation from 10 random seeds.

### Control and learning with spiking feedback

So far we have considered a continuous feedback signal, representing diffuse transmission of a neuromodulator. But many important feedback pathways in the brain communicate via spikes; we now generalize the theory to incorporate these signals. Our modeling choices are specifically motivated by the sparse climbing-fiber inputs to cerebellar Purkinje cells, which provide instructive signals that drive plasticity at parallel-fiber synapses (Hull and Regehr 2022; Silva et al. 2024). However, the general formulation is applicable to other scenarios, such as dendritic spikes in cortical neurons that are triggered by feedback connections (Larkum 2013; Richards and Lillicrap 2019; Fişek et al. 2023).

We model the spiking feedback signal, *f*_*S*_, as an inhomogeneous Poisson process with rate

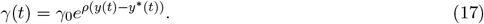

Concretely, *f*_*S*_ comprises a series of delta functions centered at spike times that are generated with instantaneous rate *γ*(*t*). The synapses ‘see’ the spikes, and can use those to update synaptic strength. The parameters *γ*_0_ and *ρ* set the spontaneous rate of feedback and steepness of the nonlinear response to output error (representing, for instance, the transfer function of an error-coding neuron).

With spiking feedback, the standard formalism cannot be used for estimating the error and target weights. Instead, similar to Eden et al. (2004) and Pfister et al. (2010), we derived a recursive Bayesian algorithm. The details are provided in *Methods: Spiking feedback*, with the full plasticity rule described by equation (117). This has a similar structure to the rule discussed above (equations (13) and (16)). Incoming feedback spikes are optimally filtered to form an estimate, 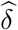, of the continuous-valued output error (Fig. 3a). This is used to set the fast weights via an optimal control gain as before. For the slow weights, 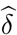 is transformed into an estimate of the instantaneous feedback rate, 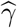, and combined with an eligibility trace, *z*_*i*_, to drive learning,

**Figure 3:**
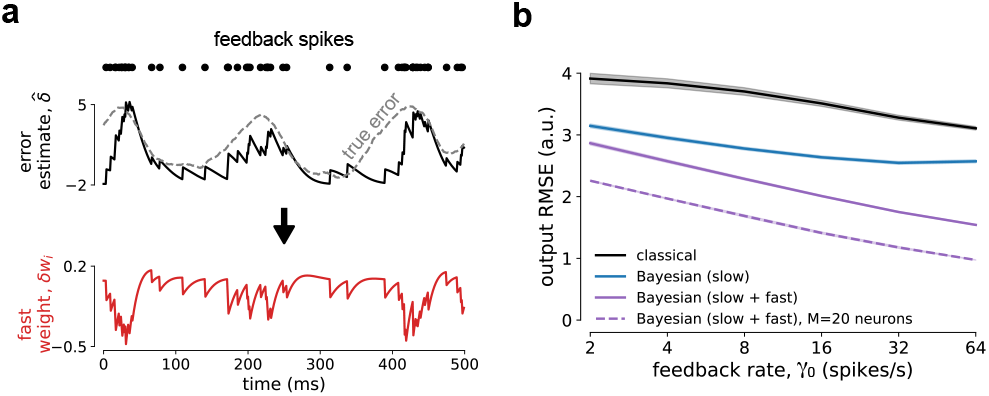
Control and learning with spiking feedback. **a)** When feedback is communicated via noisy spiking activity, optimal plasticity requires the synapse to infer the true underlying continuous error signal. The estimated error is then used to drive both control and learning. Fast weights seek to improve performance by canceling the estimated error (lower panel), while slow weights seek to reduce the error permanently. **b)** Performance depends on the rate of feedback. High feedback spike rates increase the precision of error estimates, thereby enhancing learning and control. A population of neurons acting on multiple independent feedback signals (dashed purple trace) can compensate for very low individual feedback rates. Shaded areas are standard deviation from 10 random seeds.

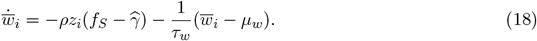

The eligibility trace, which evolves similarly to equation (16c), records the recent activity level of the synapse. Qualitatively, the slow weights of recently active synapses tend to decrease in steps when feedback spikes arrive and increase gradually in between. This process eventually reaches a steady state where the observed feedback rate is equal to the predicted rate, which happens when the target weights have been accurately estimated.

We compare the Bayesian rules to a spiking-feedback version of the classical rule used in the continuous model above (*Methods*, equation (38)). Notably, the classical spike-based rule conforms to the standard model of Purkinje cell plasticity (Ito 2001; Coesmans et al. 2004): synaptic input alone leads to long-term potentiation (LTP), whereas the conjunction of synaptic input and feedback spikes are required for long-term depression (LTD).

Simulations with the spiking-feedback model were consistent with those of the continuous model, recapitulating all of the previously observed advantages of Bayesian plasticity (Figs. 3b and S2). Learning in this case is substantially harder, requiring an order of magnitude more time to reach equivalent levels of performance, primarily because spikes occur rarely. The spontaneous feedback rate, *γ*_0_, is therefore a key determinant of performance, with higher rates leading to faster learning and lower output error (Fig. S2).

Not only does a low feedback rate make learning hard, it also makes fast-weight updates less effective. This is not surprising, since online corrections can only be reliably made soon after receiving an error-encoding spike. However, we found performance can be boosted by distributing the task and feedback across multiple neurons. We explored this by simulating *M* = 20 neurons, each as described by equation (7). For ease of comparison with the single-neuron simulations, we placed 50 synapses on each of the *M* neurons to give the same total number of synapses. The firing rates in the population model, *r*_*m*_, are summed to drive a common downstream output, modifying equation (9),

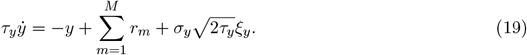

Each neuron receives a separate Poisson feedback signal, with spikes generated independently from the common error-dependent rate *γ*(*t*). Because the collected feedback spikes now provide a denser error signal, effective control can be achieved at the population level even when *γ*_0_ is small (Fig. 3b, dashed line).

### Fast and slow synaptic plasticity in the cerebellum

Our theory was derived in a setting where the target weights are defined explicitly and the target output is guaranteed to be realizable by the model. To validate our results in a biological setting and, cucially, to make experimentally testable predictions, we apply it to a cerebellar learning problem.

Our population model can be mapped onto the structure of a cerebellar microzone – a group of Purkinje cells receiving correlated climbing-fiber input and whose outputs converge on common down-stream targets (Fig. 4a) (Apps et al. 2018). As an abstraction of microzone processing, we consider a population of 20 neurons that must learn to transform patterns of synaptic input into time-varying outputs (Fig. 4b). The inputs come from parallel fiber spikes, assumed to be driven by time-varying patterns of granule cell activity (Medina et al. 2000; Gilmer et al. 2023). The output represents a downstream motor signal, or a predicted motor output (Wolpert et al. 1998). Feedback is provided by sparse climbing-fiber spikes, with *γ*_0_ = 2 spikes/s, leading to firing rates close to that observed *in vivo* (Armstrong and Rawson 1979).

**Figure 4:**
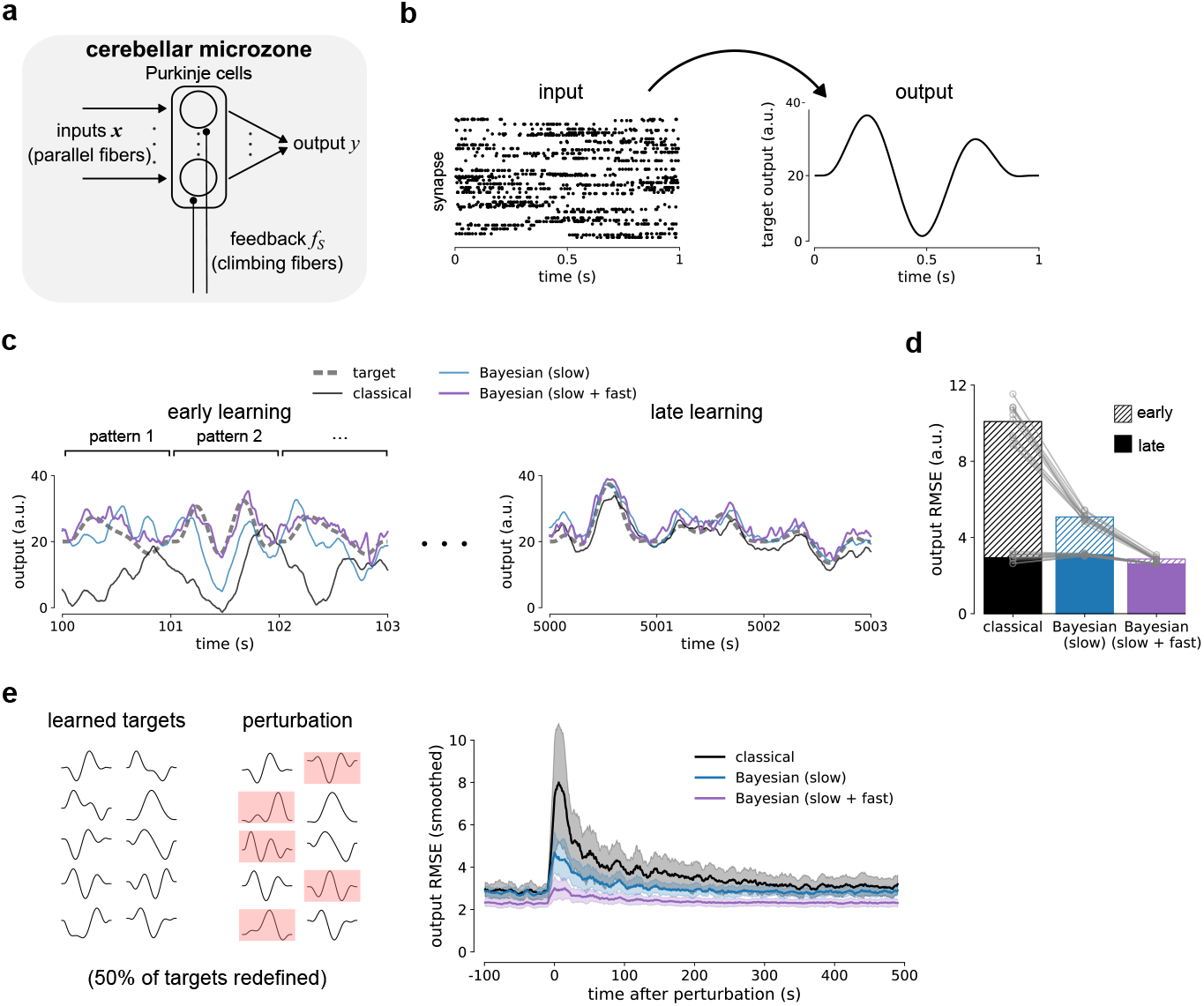
Fast and slow plasticity in the cerebellum. **a)** Schematic of the cerebellar microzone model. A group of Purkinje cells receiving synaptic input from parallel fibers project to a common downstream output. Each Purkinje cell receives feedback spikes from a separate climbing fiber, signaling the error between actual and target outputs. **b)** Schematic of the learning task. Ten patterns of temporally organized parallel-fiber input must be mapped by a population of Purkinje cells to time-varying target outputs. **c)** Example outputs early and late in learning, replicating the results from the toy model in Fig. 1b. **d)** Performance during early (100 − 200 s) and late (5000 − 5100 s) stages of learning. Bars denote RMSE between output and target output, averaged over 100 s. **e)** Left: After training on 10 input-output mappings, half of the target outputs were changed. Right: Bayesian plasticity confers faster recovery than the classical approach. With fast weights, the output is largely insensitive to the perturbation. Error curves have been smoothed with a moving average filter of width 10 s. Shaded areas are standard deviations from simulations pooled over 10 random training seeds × 10 perturbations.

We defined 10 pairs of input-output patterns {***ν***_*p*_(*t*), 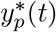; *p* = 1, …, 10}, each lasting 1 s (Fig. 4b). The input patterns comprised time-varying vectors of synaptic input rates, ***ν***_*p*_, realized with Poisson spiking. We used 2000 synapses in total, distributed equally across the 20 Purkinje cells. The target output trajectories were defined as random wavelets with shapes that drift on a slow timescale of ∼10^4^ s (analogous to the target-weight drift in the teacher-student task). Input-output pairs were presented in a continuous stream in random order for 500 s. While all of the models can eventually solve this task, the Bayesian models learn much faster, and control via the fast weights can pin the output to the target from the outset (Fig. 4c,d).

So far we have focused on learning a particular input-output mapping. What happens when the desired input-output mapping changes? To address this, we tested the response of the trained models to changes in the target outputs. As above, we trained on 10 input-output mappings, but after training we changed 5 of the outputs (Fig. 4e). An efficient response requires synapses to adapt selectively to learn the new input-output pairs, without disrupting previously learned pairs. We find that Bayesian plasticity is much more robust than the classical approach, enabling rapid recovery over a few tens of pattern presentations. With fast-weight updates, the output error is almost unaffected by the perturbation. And in the background, the slow weights gradually converge to their new target values, reducing the magnitude of ongoing control currents and updating the stored memory of the task.

### Signatures of fast and slow synaptic plasticity

We use the cerebellar learning task to make experimental predictions for the control and learning components of our theory. In the cerebellar model, the control signal depends strongly on climbing-fiber input, transiently adjusting activity whenever a feedback spike arrives. To measure the size of this effect in our model, we plotted the firing rates of neurons relative to the time of their associated climbing-fiber spikes. These simulations show a pronounced dip in firing rates after a climbing-fiber spike, but only when there are fast-weight updates; the dip disappears both for classical and Bayesian updates of slow weights only (Fig. 5a). Such patterns of activity – climbing-fiber-induced ‘spike pauses’ – have been widely observed *in vivo* (Bell and Grimm 1969; Bloedel and Roberts 1971; Sato et al. 1992; Barmack and Yakhnitsa 2003; Han et al. 2020). However, while widely observed, their computational significance has remained unclear; our model provides a novel, normative explanation that spike pauses are a strategy for suppressing downstream error. Moreover, it makes two experimentally testable predictions: the duration of the dip increases with the time constant of downstream dynamics, *τ*_*y*_, and the magnitude of the dip decreases with feedback delays (Fig. 5b). Both predictions could be tested by manipulating output dynamics experimentally, or exploiting the fact that feedback delays vary across different regions of the cerebellum (Suvrathan et al. 2016; Jayabal et al. 2022).

**Figure 5:**
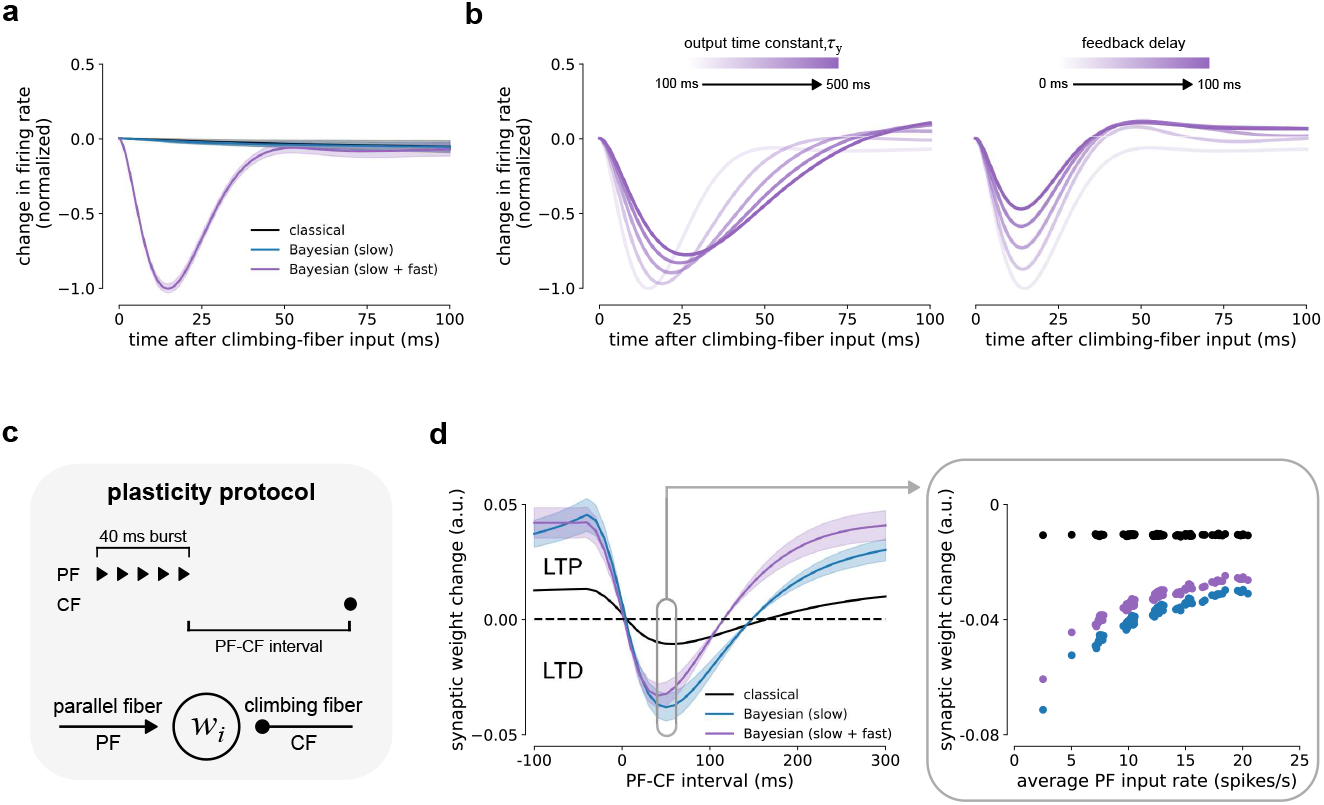
Signatures of fast and slow synaptic plasticity. **a)** Suppression of Purkinje-cell firing rates after climbing-fiber input is a signature of fast-weight updates. The solid lines denote averages of firing rates aligned to climbing-fiber feedback spikes during the cerebellar learning task. Shaded areas are the standard deviation of the average firing rate change of 20 neurons × 10 random seeds. **b)** The duration and magnitude of the predicted firing rate suppression vary systematically with the time constant of the downstream output and the feedback delay. **c)** A standard experimental protocol can discriminate between classical and Bayesian plasticity. After training on the cerebellar learning task, synapses were stimulated with conjunctions of parallel-fiber input bursts and climbing-fiber feedback spikes (50 repetitions at 2 reps/s). **d)** Left: The Bayesian rules produce LTD over a narrower range of PF-CF intervals than the classical rule, and also exhibit greater variability across synapses. Right: The variability is due to the adaptive learning rate: in the Bayesian rules, unlike the classical rule, the magnitude of weight changes depend on the average rate of input during the task. Solid lines denote synaptic weight changes measured at the end of the protocol, averaged over 100 tested synapses. Shaded areas are standard deviation.

Our model also makes predictions about the slow-weight dynamics. For that we used the trained models to simulate a common experimental plasticity protocol. In this protocol, synapses are repeatedly stimulated with a short burst of parallel fiber spikes, followed by one or more climbing-fiber spikes (here repeated 50 times at 2 repetitions/s) (Fig. 5c). In experiments, LTP is generally observed at zero or negative intervals between parallel and climbing-fiber stimulation, but switches to LTD for sufficiently positive intervals or sufficiently many climbing-fiber spikes (Wang et al. 2000; Safo and Regehr 2008; Mathy et al. 2009; Suvrathan et al. 2016; Bouvier et al. 2018). All of our models reproduced these observations, and have qualitatively similar behavior as the parameters of the plasticity protocol are varied (Fig. S3). However, the Bayesian rules yielded sharper tuning to the interval, and, importantly, greater variability across synapses (Fig. 5d). The source of this variability is the adaptive learning rate of Bayesian plasticity: the magnitude of slow-weight changes during the protocol depends on the average rate of input received during the task. Our theory can therefore be discriminated from the standard model by combining well-established plasticity protocols with recording or manipulation of input rates. While this may be possible *in vivo* in the near future with optogenetics and voltage imaging (Fan et al. 2023), *in vitro*, we predict that preconditioning a synapse with a long train of parallel fiber spikes can block subsequent plasticity induction by driving down its intrinsic learning rate.

## Discussion

We proposed that synaptic plasticity should operate at two timescales: a fast timescale to suppress immediate errors, and a slow timescale to learn. This multiscale plasticity rule significantly outperforms online gradient-based learning, confers robustness to noise and other perturbations, and means that neurons can accurately perform a task almost immediately – well before learning has stored a longer-term solution in the weights.

### Fast weights for feedback control

Our theory frames synaptic plasticity as an optimal control problem. This is a natural framework for any system where real-time feedback is available to guide the dynamics. While there is a long tradition of using control theory to understand neural function (Todorov 2004; McNamee and Wolpert 2019), our work is most closely related to recent studies that have also connected fast control mechanisms with slower processes of learning. In one approach, fast feedback was used to drive spiking network dynamics to a regime that enables local learning of complex tasks (Bourdoukan and Denève 2015; Denève et al. 2017; Alemi et al. 2018). In another, the Deep Feedback Control theory uses feedback to drive network output to a target steady-state, and then exploits credit assignment signals implicit within the controller to gradually tune feedforward weights (Meulemans et al. 2021; Meulemans et al. 2022; Rossbroich and Zenke 2023). In a recurrent network model of motor adaptation, Feulner et al. (2022) notably used the same error feedback signal for both control and learning, finding network dynamics resembling data from monkeys. Our theory differs markedly from all of these approaches, however, as the control signals in our models can be computed and implemented by individual synapses. Thus, in place of complex network-level control structures, we are positing a novel processing role for intracellular signaling at the synapse (Bhalla 2014; Benna and Fusi 2016; Zenke et al. 2017). Exploiting this local layer of processing could allow the brain to operate much more efficiently: physically, it reduces the amount of wiring and neurons needed for high performance; algorithmically, neurons make dual use of the feedback signals they receive.

The fast mechanism we propose differs from typical forms of short-term plasticity (Zucker and Regehr 2002; Abbott and Regehr 2004). Namely, in our model, fast weight changes depend strongly on error feedback, rather than presynaptic spike patterns alone. How could this be implemented biologically? If error signals are carried by a neuromodulator, as assumed in our continuous-feedback model, there are a range of candidate pathways. Neuromodulators including acetylcholine (McGehee et al. 1995; Gil et al. 1997), serotonin (Feng et al. 2001), dopamine (Higley and Sabatini 2010), norepinephrine (Cheun and Yeh 1992), among others (Marder 2012; Nadim and Bucher 2014), are all capable of selectively regulating synaptic transmission and the magnitudes of postsynaptic currents. In common with our model, these effects are transient, can be bidirectional, and are distinct from influences on membrane excitability and long-term plasticity. Alternatively, feedback signals might be processed by closely apposed glia. For instance, astrocytes would be well-placed to perform the computations we describe: they are known to integrate a variety of molecular signals and dynamically fine-tune both both pre and postsynaptic properties (Perea et al. 2009; De Pittà et al. 2016; Papouin et al. 2017).

In our cerebellar model, we assume that the fast fluctuations are driven by climbing fiber input. At a functional level, decades of past experiments are in striking agreement with our theory: when a climbing fiber spike is received by a Purkinje cell, there is a brief pause in output firing. Previous functional explanations for spike pauses include precise temporal or multiplexed coding (Steuber et al. 2007; De Schutter and Steuber 2009; Han et al. 2020), encoding dendritic spike rates (Davie et al. 2008), and transmitting teaching signals to downstream neurons (Mathews et al. 2012). While not mutually exclusive, we offer the simple explanation that if climbing fibers are indeed signaling error, then pauses are an effective mechanism for making short-term corrections. Our modeling predicts this mechanism would be most apparent in experiments focused on regions and modalities with short feedback delays, such as those receiving signals from the spinal cord for postural maintenance, or proprioceptive feedback from the periphery. It will be interesting to investigate how feedback control of cerebellar output could also support the function of more complex, nested motor-control loops (Wolpert et al. 1998; Rotondo et al. 2023), and cognitive computations more generally (Hull 2020; Kostadinov and Häusser 2022; Pemberton et al. 2023).

### Slow weights for learning

Optimizing weights for control on fast timescales also leads to highly efficient learning on slow timescales. The rule that arises naturally in our framework works efficiently by maintaining a Bayesian estimate of the target synaptic weight and its uncertainty – the best a synapse can do, given a sequence of noisy local observations. Updating weights slowly in line with this evolving estimate gradually reduces the magnitude of corrections that need to be applied via the fast weights.

Bayesian approaches to plasticity were explored in much earlier work, both in machine learning (Buntine and Weigend 1991; MacKay 1992) and neuroscience (Dayan and Kakade 2000), but have only recently begun to be broadly exploited (Blundell et al. 2015; Hernández-Lobato and Adams 2015; Kappel et al. 2015; Kirkpatrick et al. 2017; Drugowitsch et al. 2019; Hiratani and Latham 2020; Aitchison et al. 2021; Jegminat et al. 2022; Malkin et al. 2024). Aitchison et al. (2021), in particular, were the first to develop a Bayesian theory of local synaptic plasticity. In common with our slow-weight rule, they found that tracking weight uncertainty leads to an adaptive learning rate, yielding superior performance to a simple online delta rule. As well as the novel connection to control, the slow-weight component of our theory generalizes their results in two important directions. First, Aitchison et al. (2021) assume that data seen by a synapse comprises independent samples, approximating away any history dependence. By contrast, our dynamic model and learning rule accounts for, and crucially exploits, the temporal correlations inherent in neural signals. Second, our theory incorporates sources of spiking feedback and communication delays, making Bayesian plasticity more widely applicable as a theory of learning in the brain. Simulating our more realistic model, we outlined how this theory could be tested (Fig. 5): if synapses have evolved to use local data as efficiently as possible, then adding a pre-conditioning train of input to common *in vitro* plasticity protocols should unmask the characteristic adaptive learning rate.

### Outlook

Our general framework can be used to make predictions about plasticity in other cells and circuits, including more complex scenarios than we have studied here. For instance, while we have focused on linear neural dynamics – a suitable approximation for cerebellar Purkinje cells (Llinás and Sugimori 1980; Brunel et al. 2004; Walter and Khodakhah 2006) – the theory could be adapted to account for nonlinear integration in cortical and hippocampal neurons (Poirazi et al. 2003; Payeur et al. 2021; Bicknell and Häusser 2021). This could be achieved with a straightforward modification of the Kalman filtering approach that we used, for which there are already established theoretical tools for nonlinear filtering and control (Crassidis and Junkins 2011; Kutschireiter et al. 2020). At the network level, combining fast and slow plasticity with recurrent connectivity would be a challenging, but likely very fruitful, direction. It has previously been shown, for instance, that the concerted action of multiple plasticity rules can enhance memory formation and stability (Zenke et al. 2015).

Finally, to derive our fast and slow plasticity rules, the synapses had to know the underlying equations that transform synaptic drive to the output. How this is learned is an interesting, and important, problem. It is likely that some of the learning can be accomplished on much slower, even evolutionary, timescales (Friedrich et al. 2021). On shorter timescales, Moore et al. (2024) proposed an elegant solution: learn the optimal controller from data without ever learning the underlying equations. Applying this formalism, they demonstrated that a wide range of neurophysiological data can be explained by modeling individual neurons as optimal feedback controllers. Incorporating similar ideas to study nonlinear computations in hierarchical and recurrent neural circuits is an important avenue for future work.

## Methods

### Online linear regression

Using the simple linear regression model described by equation (1), we illustrated how concurrent fast and slow updates to synaptic weights can be exploited to optimize task performance. Here we provide the derivation of the plasticity rules for that model (equations (4) and (6)).

We consider a discrete-time setting where the inputs, output and weights are updated at small intervals, Δ*t*. Decomposing synaptic weights into fast and slow components, equation (1) becomes

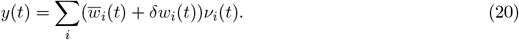

After observing the local inputs, *ν*_*i*_, and error, *y* − *y*\*, at time *t*, synapses update their fast and slow weight components in parallel,

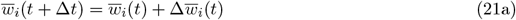

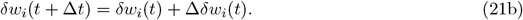

The role of the fast weights, *δw*_*i*_, is to greedily suppress the current error, so that *y* approximately matches *y*\* – independent of the setting of the slow weights. We achieve this by choosing the fast-weight updates to be proportional to the negative of the error,

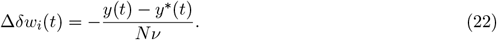

where *N* is the total number of synapses and 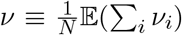 is the expected input rate. Using this expression, and inserting equation (21) into equation (20), at the next time step we have

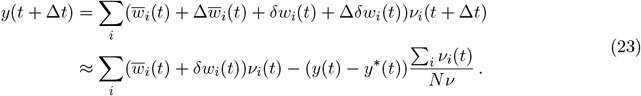

To derive the approximate expression in the second line, we replaced *ν*_*i*_(*t* +Δ*t*) by *ν*_*i*_(*t*), which is valid if Δ*t* is small compared to the timescale over which the firing rates change, and dropped the term 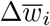, which is valid if the slow weight updates are small. The first term is just *y*(*t*) (see equation (20)); approximating Σ_*i*_ *ν*_*i*_ by its expectation then gives

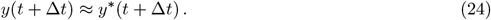

Thus, with mild assumptions, the fast-weight updates given in equation (22) force the output to closely match the target at all times.

While the fast-weight updates ensure low output error, this comes at the cost of ongoing, and potentially large, fluctuations in weights. Those fluctuations can be seen in the right panel of Fig. 1a, especially at early times before the slow weights have been updated. We address this with a local rule for the slow weights that permanently pushes *y* closer to *y*\*, thereby reducing the need for ongoing corrections. To that end, at each time step, we adjust the slow weights with the aim of minimizing the loss

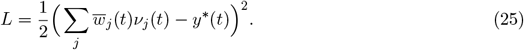

The loss is minimized by making updates in steps proportional to the negative of the gradient

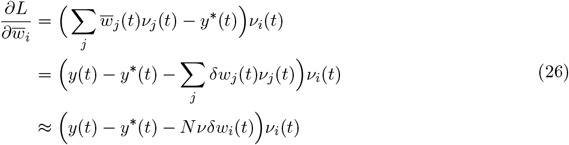

For the last line we used the fact that the fast weights do not depend on the synapse index, so *Σ*_*j*_ *δw*_*j*_*ν*_*j*_ = *δw*_*i*_ *Σ*_*j*_ *ν*_*j*_ ≈ *Nνδw*_*i*_. This leads to a local slow-weight update rule

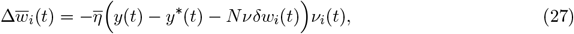

where 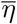 is a small learning rate. Equations (22) and (27) for the fast and slow updates correspond to equations (4) and (6) in the main text.

### Classical online learning

For the model described by equations (7) and (9), with either continuous feedback (equation (10)) or spiking feedback (equation (17)), we compared our Bayesian plasticity rules to classical, gradient-based counterparts. The classical rules are based on the Real Time Recurrent Learning algorithm (RTRL; Williams and Zipser 1989; Pearlmutter 1995), which is the canonical online, gradient-based learning rule for dynamic models. While typically employed in recurrent neural networks, here we apply it to a single-neuron model, adapted slightly to enable fair comparisons with alternatives.

For continuous feedback, we aim to minimize the squared output error, (*y* − *y*\*)^2^. However, unlike the simple model, here we need to account for history dependence – the output at any given time depends on inputs that were received in the past. To account for this, we start by considering a loss integrated over a time horizon, *h*, in which the weights are held fixed,

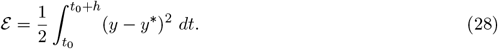

The derivative of the loss with respect to the weight of synapse *i* over that period is

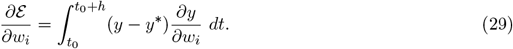

The second factor in the integral is a dynamic quantity, which can be computed by taking derivatives of equations (7) and (9) with respect to *w*_*i*_. This gives the linear system

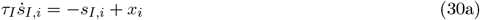

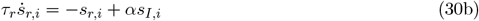

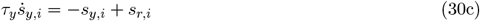

for the sensitivities 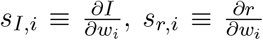, and 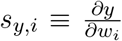. Equation (30) is solved from the initial condition *s*_*I,i*_(*t*_0_), *s*_*I,i*_(*t*_0_), *s*_*I,i*_(*t*_0_) = 0. An exact gradient-descent weight update could then be applied at time *t*_0_ + *h* as

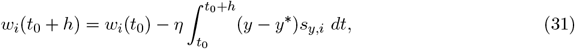

where *η* is the learning rate. Functionally, *s*_*y,i*_ plays the role of an eligibility trace by recording the recent impact of the synapse on the output.

Formally, after the weights have been updated, the sensitivities should then be reset to the zero initial condition to track activity over the next time interval. However, assuming weight changes are slow compared to the model dynamics, the RTRL rule is commonly run as a fully online approximation by taking *h* → 0, giving a differential equation for the weight dynamics, and neglecting the reset of the sensitivities (Pearlmutter 1995). We take this approach here and make two further modifications. First, we replace the term (*y* − *y*\*) with its noisy observation, *f*. Second, we append an additional weight decay term to account for the slow drift of the target weights (see Aitchison et al. (2021) and the derivation of the Bayesian rule below for motivation). This leads to classical online updates for each synapse

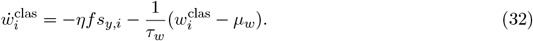

For spiking feedback, we use the same approach, but with a different loss function. In this case the output error is communicated via a Poisson process with rate *γ*(*t*), as described by equation (17). To handle the random and discrete nature of spiking feedback, we work with spike probabilities.

Over a time horizon *h*, the feedback spike count, denoted *n*, will be distributed as

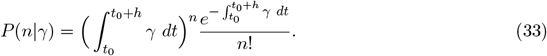

Updates to the weights should seek to match this distribution to the one that would arise if all of the weights were set correctly. We express this in terms of a ‘target rate’, *γ*\*, which defines a target distribution, *P* (*n*|*γ*\*), via equation (33).

We use the cross-entropy loss between target and actual distributions to derive the updates; this is given by

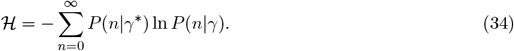

The derivative of the loss with respect to weight *w*_*i*_ is, using 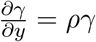,

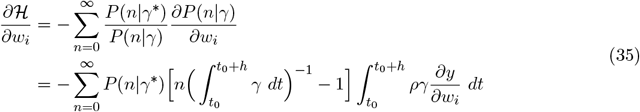

As before, we use equation (30) to compute the 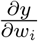 term, accounting for the history dependence.

Similar to the continuous feedback model, we define weight updates to be proportional to the negative of the loss gradient, and then take *h* → 0 to yield a continJuous-time expression. Expanding equation (35) to first order in *h*, approximating the integrals as 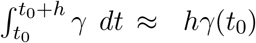, and exponentials as *e*^−*hγ*^ ≈ 1 − *hγ*, only the *n* = 0 and *n* = 1 terms survive from the sum. Some straightforward algebra then leads to

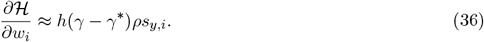

Thus, the spiking-feedback analogue of equation (31) is

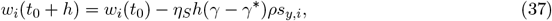

where *η*_*S*_ is the learning rate.

Taking *h* → 0, and approximating the unobserved rate *γ* with feedback spikes, *f*_*S*_, leads to a continuous-time learning rule

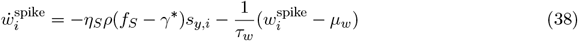

where we have appended a weight decay term as before.

Finally, to set *γ*\*, we use the fact that when the weights are set correctly, the error comprises Gaussian fluctuations about zero due to noise. With the exponential nonlinearity in the feedback rate function (equation (17)), the expected rate is the mean of a log-normal variable, giving

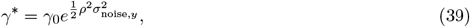

where 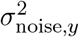 is the stationary variance of the output noise. The stationary variance can be computed numerically in terms of the model parameters via the Lyapunov equation

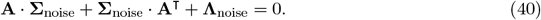

In equation (40), **A** is a matrix encoding the model dynamics (see equation (70), below), and **Λ**_noise_ is diagonal matrix comprising noise variances 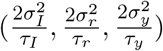 on the diagonal. The relevant term 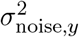 is found at the last row and last column of **Σ**_noise_.

In all simulations, learning rates for the classical rules were selected via grid search to minimize output error.

### Derivation of the Bayesian plasticity rule

#### Problem setting

Given the model dynamics described by equations (7) and (9), we consider a general task in which the goal is choose weights, *w*_*i*_, to match the output, *y*, to a time-varying target, *y*\*. We assume *y*\* is generated via deterministic dynamics identical to *y*, using a set of target weights 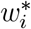. Following Aitchison et al. (2021), to represent environmental variability or ongoing changes in the local circuit, we assume the mapping between input and target output drifts slowly over time, modeled as random drift of the target weights about a constant mean, *µ*_*w*_,

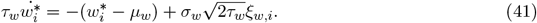

The drift timescale, *τ*_*w*_ ∼ 1000 s, is orders of magnitude slower than the neuronal and output dynamics.

The above assumptions about the relationship of target weights to *y*\*, and the drift described by equation (41), are exact in the teacher-student learning paradigm. In the cerebellar learning simulations, target weights are instead defined implicitly as weight values that would solve the task.

#### Error minimization from the perspective of a single synapse

As in the simple online linear regression model, we decompose the weights into independent slow and fast components, 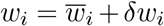. The fast weights, *δw*_*i*_, will be responsible for suppressing error, while the slow weights, 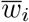, gradually converge to the target weights, thereby reducing the magnitude of ongoing corrections. However, with the more realistic dynamics described by equations (7) and (9), writing down the plasticity rule is considerably more challenging. That’s because the mapping from the weights, *w*_*i*_, to the output, *y*, is more complicated: rather than a simple sum, as in equation (1), there are several differential equations separating the weights from the output (equations (7) and (9)).

To derive learning rules in this more complicated setting, we start by introducing a set of variables equal to the difference between actual and target quantities,

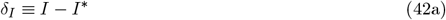

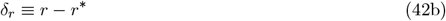

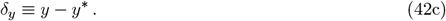

From equations (7) and (9), these evolve via

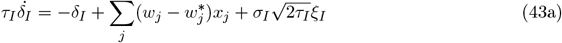

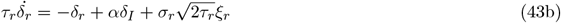

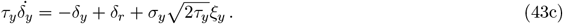

The target weights, 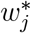, are unknown, so on the surface these equations don’t seem especially useful. However, the synapses have access to the feedback, *f*, which is a noisy version of *δ*_*y*_ (see equation (10)). Thus, they can use these equations to estimate 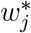 based on the mismatch between *δ*_*y*_ and *f*, and update the actual weights, *w*_*j*_, with the aim of making *δ*_*y*_ as small as possible.

However, there’s a problem: *δ*_*y*_ depends on the activity of all the synapses; information that any one synapse doesn’t have. To remedy this, we’ll work from the perspective of synapse *i*, and model *δ*_*y*_ in terms of local variables.

Decomposing synaptic weights into slow and fast components, the sum in equation (43a) can be expanded as

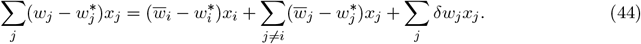

The first term depends only on synapse *i*, so it’s local. The second term represents a source of error due to the other *N* − 1 unobserved synapses. For Poisson input and large *N*, this can be approximated as Gaussian white noise (Fourcaud and Brunel 2002),

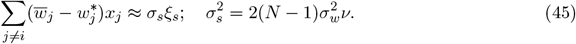

As above, *ν* denotes the expected input rate per synapse.

For the third term in equation (44), we view the sum over fast weights as approximating a single scalar control variable, denoted *u*, later to be chosen to minimize output error. Assuming that all of the synapses can compute *u* locally, by setting the fast weights uniformly as

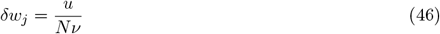

the third term in equation (44) can be expressed as

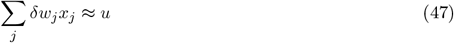

where the approximation is good in the large *N* limit. Inserting equations (45) and (47) into equation (44), and inserting that into equation (43), we arrive at a set of equations that uses only information local to synapse *i*,

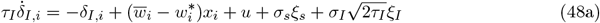

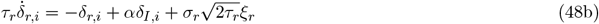

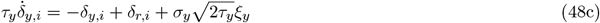

Equation (48) is a model of the error that the synapse can use to guide plasticity. In our rule, the synapse learns by updating its slow weight, 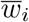, to be equal to an evolving estimate of the target, 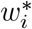, while choosing *u*, and thereby the fast weight, *δw*_*i*_, to cancel out the remaining error.

### Optimal control solution

Because the noise in equation (48) is Gaussian and we want to minimize the squared error, (*y* − *y*\*)^2^, the problem of finding the optimal *u* reduces to the well-known linear-quadratic-Gaussian (LQG) control problem (Crassidis and Junkins 2011). In this setting, the optimal control, denoted 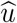, satisfies

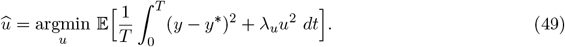

Note that this includes a control cost, *λ*_*u*_*u*^2^, which limits the size of 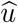; without that cost, 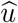 would diverge. In general, LQG control employs a Bayesian filter to estimate unobserved model variables, and then uses those estimates to construct the controller.

To solve equation (49) using standard methods, we first need to express our model for the error (equation (48)) and feedback signal (equation (10)) in a canonical linear state space form. We start by combining the model variables into a single vector,

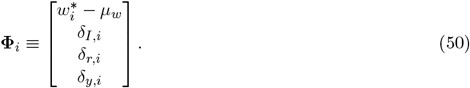

Then, we express the control variable as a vector, **u**_*i*_, and make a change of variables that will clean up an offset term,

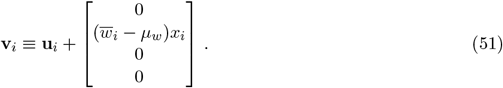

Introducing the vector

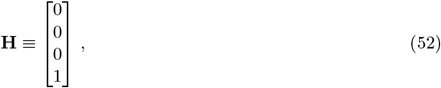

the dynamics of **Φ**_*i*_ and feedback, *f*, can be expressed together as

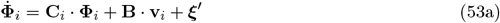

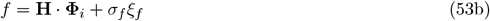

where

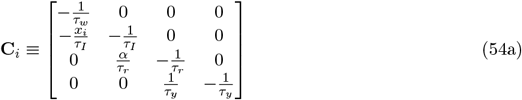

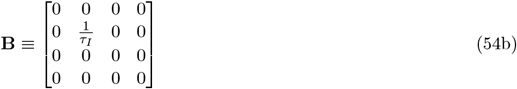

and ***ξ***′ is Gaussian white noise with covariance matrix

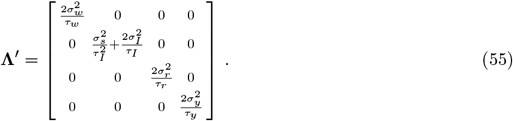

With this set up, the control problem in equation (49) can be written

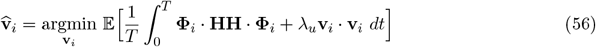

where **HH** is an outer product.

Although the control variable, **v**_*i*_, is now a vector instead of a scalar, *u*, the optimal solution will still only have one non-zero component (the second one). This is because multiplication by **B** in equation (53a) means that only the second component of **v**_*i*_ will influence the output, so the control cost term in equation (56) will force the other components to be zero. The scalar 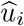 is recovered from 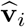 by selecting the second component and then reversing the variable change in equation (51),

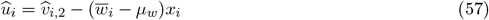

where the subscript 2 denotes the second component of the vector.

Nevertheless, equation (56) is still not identical to equation (49), because the variable change in equation (51) has shifted the quantity 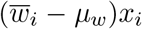 from the first term in the integral to the second term. This means *λ*_*u*_ parameterizes a slightly different control cost. We will ignore this detail, however, as it doesn’t influence any of our modeling choices or the interpretation of results.

Note also that we have written the control variable with a subscript *i*, since at this stage the solution depends on local variables. We will later make approximations that remove this dependence, so that the control variables are the same for all synapses, as assumed in equation (46).

Ignoring biological constraints, LQG theory gives the solution to the problem defined by equations (50)–(56) as

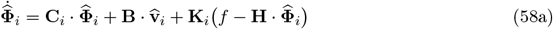

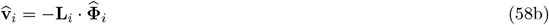

The vector 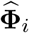 is a Bayesian (Kalman filter) estimate of the values of the unobserved variables in equation (50). **K**_*i*_ and **L**_*i*_ are optimal feedback and control gains, given by

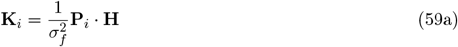

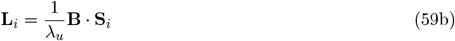

where **P**_*i*_ and **S**_*i*_ are determined by a pair of matrix Riccati equations (Crassidis and Junkins 2011),

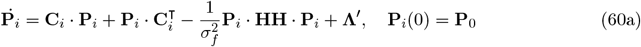

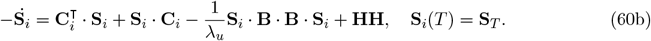

The equation for **P**_*i*_, a covariance matrix representing uncertainty about the estimate 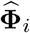, is solved forwards in time from an initial condition, whereas the equation for **S**_*i*_, used to compute the optimal control, is solved backwards in time from a terminal condition.

While these equations are exact, they are far from biologically plausible. In the next several sections we remedy that by making several approximations.

### Control gain approximation

Solving equation (60b) backwards in time poses a problem for online implementation, because the dynamics matrix, **C**_*i*_ (equation (54a)), depends on the synaptic input, *x*_*i*_, which cannot be known in advance. We therefore simply drop the term *x*_*i*_ from the matrix **C**_*i*_. With this approximation and sufficiently large *T*, equation (60b) relaxes to a steady state that is independent of the terminal condition and synapse index. The steady-state matrix **S** is then defined implicitly as the positive-semidefinite solution to the algebraic Riccati equation

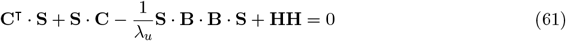

where **C** is the same as **C**_*i*_ but with *x*_*i*_ set to zero.

Writing equation (61) in component form, and using the fact that **S** is positive-semidefinite, straight-forward algebra shows that the first row and column of **S** are all zero. Consequently, because of the structure of **B** (equation (54b)), the only nonzero elements of **L**_*i*_ are the second through fourth elements of the second row. That in turn implies that only the second element of 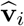 (equation (58b)) is nonzero. That element is given by

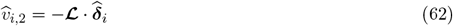

where **ℒ** is proportional to the second through fourth elements of the second row of **S**,

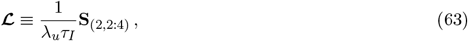

and 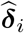 consists of the second through fourth elements of 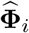,

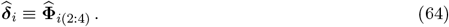

We can now use equation (58b) to derive an explicit expression for the control variable, 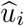,

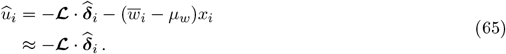

In the approximation in the last line, we ignored the second term, which will allow us to make the controller independent of *i*, and makes little difference to performance in practice.

We solve equation (61) numerically to obtain the control gain for simulations. An analytical approximation is provided in equation (91) below, for the simpler version of the plasticity rule presented in the main text; that approximation also works well in practice (Fig. S1).

### Feedback gain approximation

Although we now have all the ingredients for a local online plasticity rule, we also approximate the feedback gain **K**_*i*_ for the sake of interpretability.

We first unpack equation (60a) by partitioning the state-estimate covariance matrix into blocks,

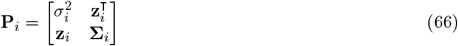

where, to highlight the functional roles that each block will play in the plasticity rule, we introduced the variables

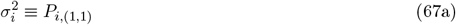

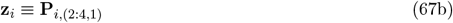

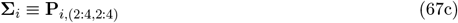

where the numeric subscripts on the right-hand side denote the range of row and column indices for each each block. The variable 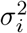 represents the uncertainty about the target weight, **z**_*i*_ represents the covariance between the target weight and error vector (see equation (64)), which will act as an eligibility trace in the plasticity rule, and **Σ**_*i*_ represents the uncertainty about the estimate of the error vector.

Defining

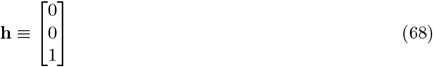

and using the definitions in equations (70) and (71), equation (60a) can be expanded to give the dynamics of each of the blocks as

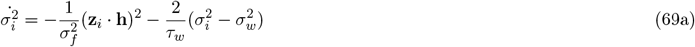

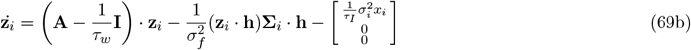

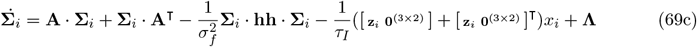

where **0**_(3*×*2)_ denotes a 3 × 2 matrix of zeros, **A** is the lower right (2 : 4) × (2 : 4) block of **C**_*i*_,

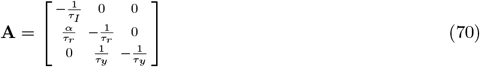

and **Λ** is the lower right (2 : 4) × (2 : 4) block of **Λ**′,

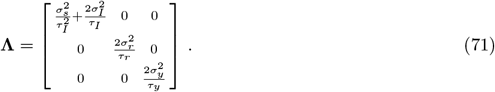

Because equations (69a) and (69b) track uncertainty related to the target weight, their temporal dynamics are strongly dependent on the local synaptic input, *x*_*i*_ – this is what that couples the weight to the observed feedback. By contrast, equation (69c) tracks the uncertainty of ***δ***_*i*_, which is dominated by uncertainty about the activity of unobserved synapses, so depends only weakly on *x*_*i*_. Mathematically, this is reflected in the fourth term of equation (69c) being dominated by the order-*N* synaptic noise term in **Λ** (see equations (45) and (71)). We can, therefore, ignore the *x*_*i*_-dependent term in equation (69c). Then, **Σ**_*i*_ relaxes to a steady state that is independent of the synapse index. That steady state is the solution to the algebraic Riccati equation,

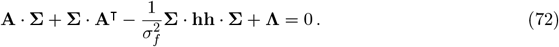

We use equation (72) in place of (69c). But to capture the most important dependence on input, *x*_*i*_, we solve equations (69a) and (69b) individually for each synapse.

Finally, we can express **K**_*i*_, equation (59a), in terms of the quantities in equation (69). Using the fact that **K**_*i*_ is proportional to the fourth column of **P**_*i*_ (see equation (52) for **H**), the first component is given by

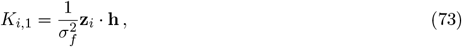

which is used to update the estimate of the target weight, 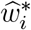. The remaining three components, used update the estimate of the error vector, 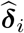, are given by the constant vector

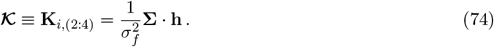

### Computations that should be performed by a synapse

Bringing everything together, we express the plasticity rule as a system of differential equations that optimally process local signals to enable concurrent control and learning.

Unpacking the error estimate from equation (58a), expressing the feedback gain via equations (72) and (74), and approximating 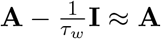, which is valid because<u>^1^</u> is orders of magnitude smaller than other terms on the diagonal of **A**, we get

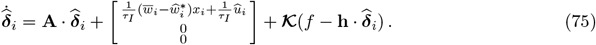

The only remaining issue is that the control variable, 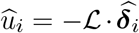 (equation (65)), still depends on the synapse index via 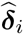. As discussed when introducing the fast weights in equation (46), we would like this to be identical for all synapses. To address this, we need only make the obvious choice of setting 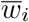 equal to its estimated target 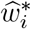 at all times. This makes the *i*-dependent term in the brackets in equation (75) vanish, and so too the *i*-dependence of 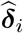 (assuming the system has been running long enough that initial conditions are forgotten).

Setting 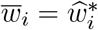, thereby dropping the synapse index on 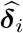 and 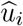, and unpacking the rest of equations (58a) and (69) leads to

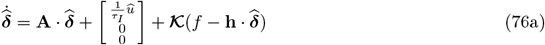

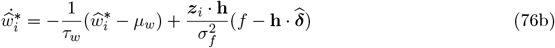

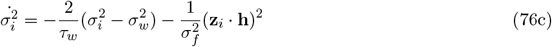

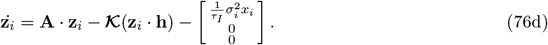

Altogether, this gives synaptic weights

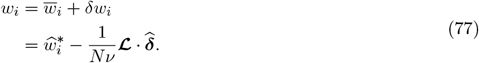

### Plasticity rule approximation for *τ*_*I*_, *τ*_*r*_ ≪ *τ*_*y*_

Control is most effective when the downstream time constant is large. This regime also permits a simple approximation of the plasticity rule that makes parameter dependencies explicit. We use these equations in the main text for ease of intuition (equations (13a) – (16b)). Here we provide the details of the approximation.

Starting from equations (76a) and (76d), we express the components of the vectors 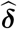 and **z**_*i*_ using *I, r* and *y* subscripts. Assuming *τ*_*I*_, *τ*_*r*_ ≪ *τ*_*y*_, and using equation (70) for **A**, we treat the *I* and *r* components as reacting instantaneously to their inputs,

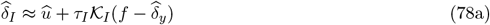

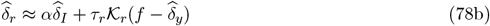

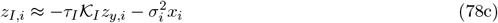

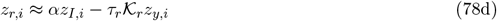

where the subscripts on 𝒦 indicate their components,

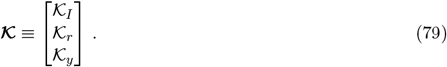

With these approximations, equation (76) becomes

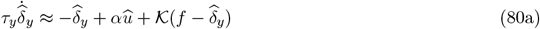

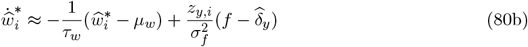

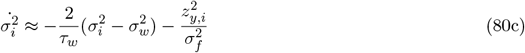

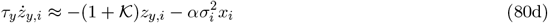

where

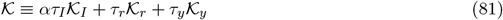

is a weighted sum of the three components of the Kalman gain (equation (74)). Equation (80) has been copied to the main-text equations (13) and (16) with some minor cosmetic changes; to clean up the presentation, there we have dropped the *y* subscripts, explicitly denoted 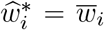, and made a sign change for the eligibility trace variable, *z*_*i*_ ≡ −*z*_*y,i*_.

To compute 𝒦, we note first of all that

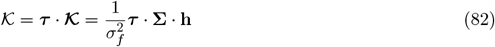

where the second equality comes from equation (74) and we have defined

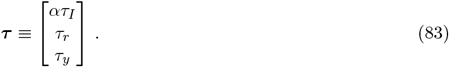

To derive an explicit expression for ***τ*** · **Σ** · **h**, we operate on both sides of equation (72) with ***τ***, giving us

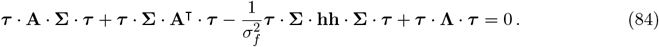

Then, combining the fact that ***τ*** · **A** = −**h** with equation (82), we arrive at

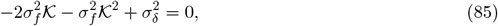

Where

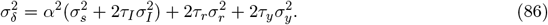

Solving equation (85) gives the scalar feedback gain explicitly as

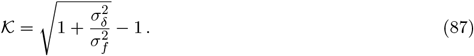

For the control gain, we make use of the duality of state-estimation and control for the LQG problem (Crassidis and Junkins 2011), which means that knowing the parameters of the equation for 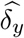 (equation (80a)) allows us to easily write down the Riccati equation needed to compute the corresponding controller. The analogue of equation (61), now for the scalar variable *s*, and with equivalent scalar parameters 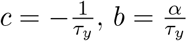 and *h* = 1, in place of **C, B** and **H**, is

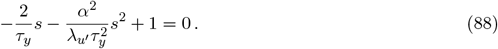

Defining

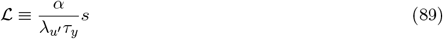

as the analogue of equation (59b) then leads to

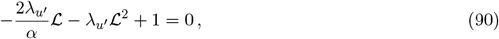

and thus a scalar control gain

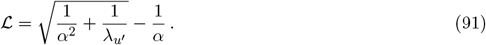

Here we have used the notation 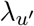 to signify that the control cost parameter is not identical to *λ*_*u*_, used above, and is fit separately in simulations of the approximate rule.

Inserting the optimal control

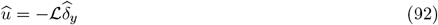

into equation (80a) yields the plasticity rule corresponding to equations (13) and (16) in the main text.

### Steady-state learning rate

Using the approximate plasticity rule given in equation (16), we can derive an expression for how the steady-state learning rate of a Bayesian synapse depends on key parameters, and especially on the firing rate.

The first observation is that because 𝒦 is typically large, after a spike *z*_*i*_ decays rapidly, and so whenever a new spike arrives it’s effectively zero. Thus, assuming for the moment that *σ*_*i*_ is constant, equation (16c) tells us that when a spike occurs at time *t* = 0, *z*_*i*_(*t*) is given by (for times *t >* 0)

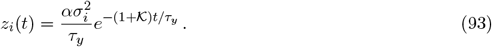

Inserting this into equations (16a) and (16b), and assuming for the moment that both are constant on a timescale of *τ*_*y*_*/𝒦*, we see that when a spike occurs they change by

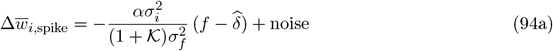

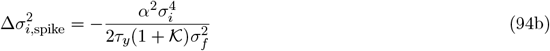

where the subscript ‘spike’ indicates that these are changes in response to a spike.

In the large 𝒦 limit, both 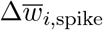 and 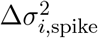 are small, justifying our assumption that *w*_*i*_ and 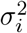 are approximately constant during a spike. The noise term in equation (94a) arises because the feedback error, *f*, has a white noise component (see equation (10)), so fluctuates around the relevant signal, *y* − *y*\*.

Because equation (94a) tells us the weight change per pre-synaptic spike, we can identify the term in front of 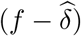 as the effective learning rate,

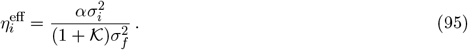

To derive an explicit expression for 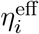, we need the steady state value of *σ*_*i*_. Using equation (16b), that’s given by

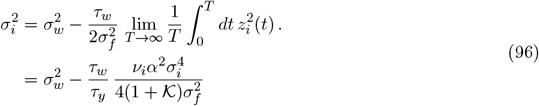

where we used equation (93) for the time course of *z*_*i*_(*t*) after a spike and, recall, *ν*_*i*_ is the input firing rate to synapse *i*. Note the slight abuse of notation: above, and in what follows in this section, 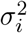 is the steady state variance; to reduce clutter we don’t make this explicit. Solving equation (96) gives us

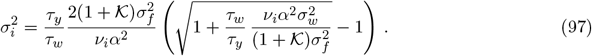

For typical parameter values (see Table 1), and using the definitions in equations (86) and (87), we find that the second term inside the square root is large (∼ 10^3^ − 10^4^) as long as input rates, *ν*_*i*_, are greater than ∼ 0.1 spikes/s. Equation (97) thus simplifies to

**Table 1:**
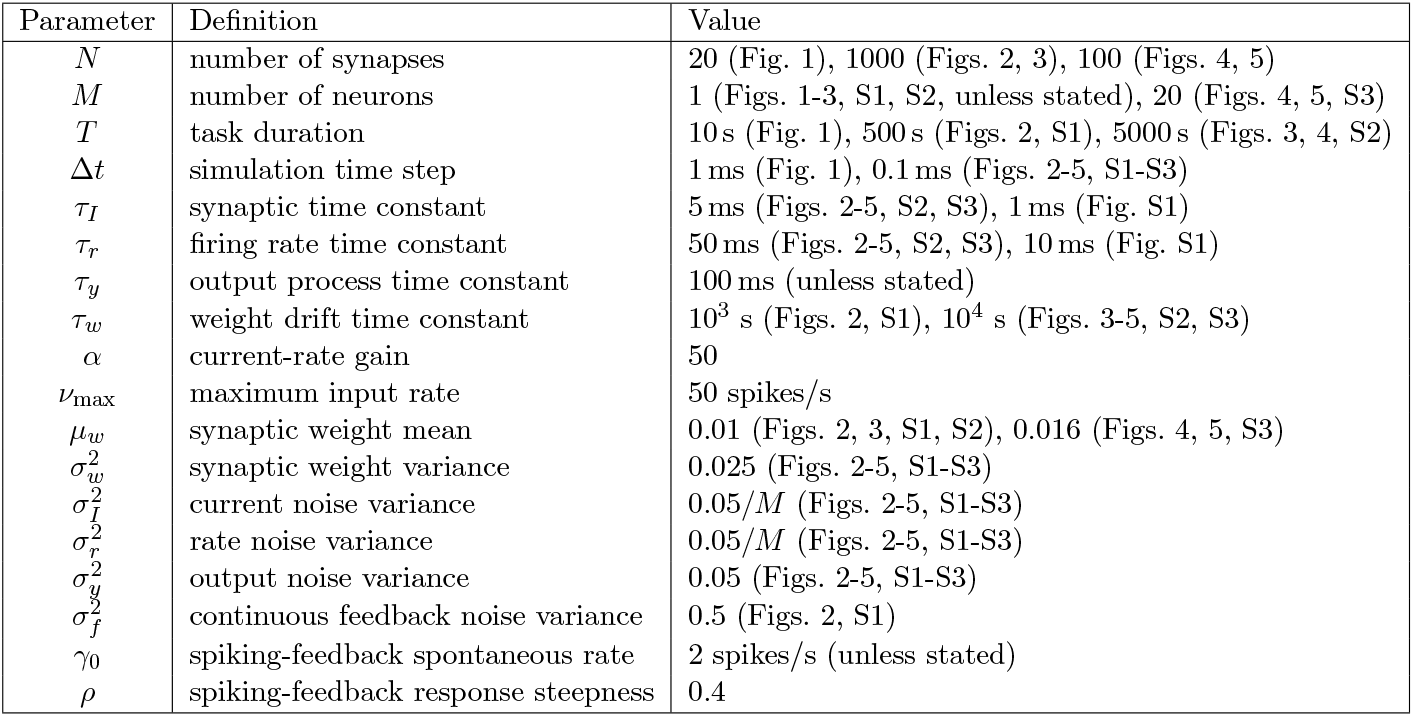
Model and simulation parameters.

| Parameter | Definition | Value |
| --- | --- | --- |
| $N$ | number of synapses | 20 (Fig. 1), 1000 (Figs. 2, 3), 100 (Figs. 4, 5) |
| $M$ | number of neurons | 1 (Figs. 1-3, S1, S2, unless stated), 20 (Figs. 4, 5, S3) |
| $T$ | task duration | 10 s (Fig. 1), 500 s (Figs. 2, S1), 5000 s (Figs. 3, 4, S2) |
| $\Delta t$ | simulation time step | 1 ms (Fig. 1), 0.1 ms (Figs. 2-5, S1-S3) |
| $\tau_I$ | synaptic time constant | 5 ms (Figs. 2-5, S2, S3), 1 ms (Fig. S1) |
| $\tau_r$ | firing rate time constant | 50 ms (Figs. 2-5, S2, S3), 10 ms (Fig. S1) |
| $\tau_y$ | output process time constant | 100 ms (unless stated) |
| $\tau_w$ | weight drift time constant | $10^3$ s (Figs. 2, S1), $10^4$ s (Figs. 3-5, S2, S3) |
| $\alpha$ | current-rate gain | 50 |
| $\nu_{\max}$ | maximum input rate | 50 spikes/s |
| $\mu_w$ | synaptic weight mean | 0.01 (Figs. 2, 3, S1, S2), 0.016 (Figs. 4, 5, S3) |
| $\sigma_w^2$ | synaptic weight variance | 0.025 (Figs. 2-5, S1-S3) |
| $\sigma_I^2$ | current noise variance | $0.05/M$ (Figs. 2-5, S1-S3) |
| $\sigma_r^2$ | rate noise variance | $0.05/M$ (Figs. 2-5, S1-S3) |
| $\sigma_y^2$ | output noise variance | 0.05 (Figs. 2-5, S1-S3) |
| $\sigma_f^2$ | continuous feedback noise variance | 0.5 (Figs. 2, S1) |
| $\gamma_0$ | spiking-feedback spontaneous rate | 2 spikes/s (unless stated) |
| $\rho$ | spiking-feedback response steepness | 0.4 |

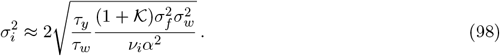

Inserting this into equation (95), we arrive at

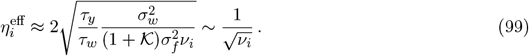

Notably, this scaling with input rate is consistent with the results of Aitchison et al. (2021), which they demonstrated was consistent with experiments.

### Spiking feedback

We now consider the case where feedback is communicated via spikes. The spiking feedback signal, *f*_*S*_, is modeled as an inhomogeneous Poisson processes with rate *γ*, as described by equation (17). The derivation is essentially the same as for the continuous feedback case above; the difference is that it’s not a textbook problem, so we have to do it ourselves. The approach, though, is relatively standard: we compute the time evolution of the mean and covariance of **Φ**_*i*_ conditioned on feedback spikes. These correspond to 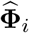 and **P**_*i*_ above, but their time evolution is, of course, different because of the spiking feedback.

### Model discretization and notation

To model feedback spikes, we discretize time into bins of size Δ*t*, and assume that there are either zero or one spike in each bin; this assumption is valid because we will eventually take the limit Δ*t* → 0. Our first step is to turn equation (53a) into discrete time updates rather than differential equations.

Discrete time variables are distinguished from their continuous versions with a *t* subscript, and several quantities are rescaled by the time step to reduce clutter. We define

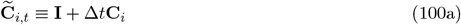

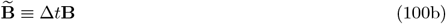

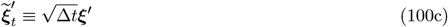

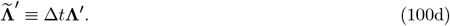

The synaptic inputs *x*_*i,t*_ are implemented as discrete-time delta functions, taking the value 1*/*Δ*t* when there is a spike and zero otherwise. The discrete-time version of equation (53a) is, then, given by

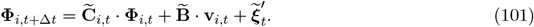

For Poisson feedback with the rate determined by equation (17), the analogue of equation (53b) is an expression for the probability of observing a feedback spike given **Φ**_*i*_. Assuming small Δ*t*, and treating spikes as Bernoulli random variables *f*_*S,t*_ ∈ {0, 1}, this is given by

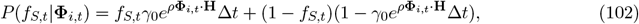

with **H** = (0, 0, 0, 1), as above (see equation (52)).

Using equations (101) and (102) as the model for feedback observations, we aim to infer a distribution over the unobserved states, **Φ**_*i,t*_, conditioned on the history of local data seen by the synapse. The data is denoted

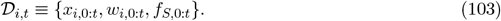

The distribution is modeled as a Gaussian with conditional mean and covariance

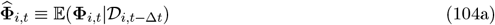

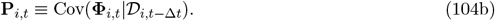

### Recursive Bayesian filter

We use standard Bayesian recursion to derive explicit expressions for the mean and covariance of **Φ**_*i*_. Our approach is similar to that of Eden et al. (2004) and Pfister et al. (2010).

At each time step, we first update the distribution over **Φ**_*i,t*_ using observation *f*_*S,t*_, via Bayes theorem, and then predict **Φ**_*i,t*+Δ*t*_ from the model dynamics,

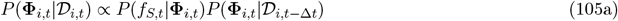

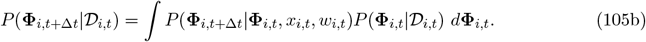

The first equation holds for our model because *f*_*S,t*_ only depends on **Φ**_*i,t*_, and **Φ**_*i,t*_ only depends on the input and weight from previous time steps. We use assumed density filtering for the update step, approximating the left-hand side of equation (105a) as Gaussian by matching the mean and covariance of the right-hand side.

When *f*_*S,t*_ = 1, the approximation is exact, giving a Gaussian

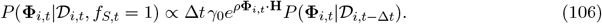

The normalizer is given by

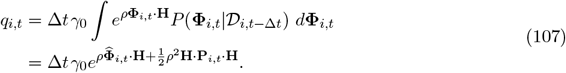

After some straightforward algebra, the updated mean and covariance are found to be

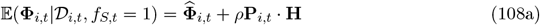

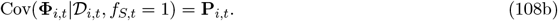

When *f*_*S,t*_ = 0,

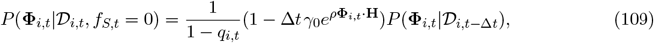

which leads to, again after some straightforward algebra,

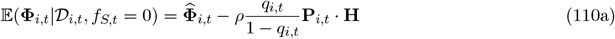

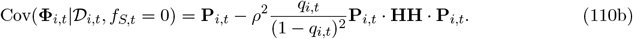

Putting both cases together, we arrive at

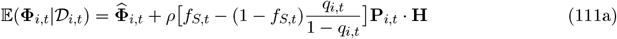

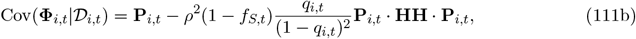

which characterizes the posterior distribution on the left-hand side of the update step in equation (105a).

Next, having completed the update step, we perform the prediction step in equation (105b) by evaluating the integral. The second factor in the integral is Gaussian by assumption, following from the previous update. The first factor, describing the state transition probability, is also Gaussian, following from the dynamics of equation (53a). This means the left-hand side of equation (105b) is Gaussian with

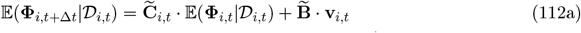

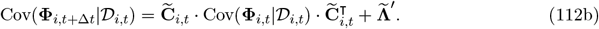

Substituting equations (111a) and (111b) into these expressions, and using the definitions in equations (104a) and (104b), gives a discrete-time estimator

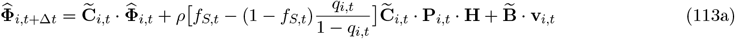

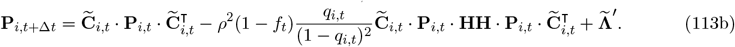

Expanding to first order in Δ*t*, using equations (100a)–(100d), and taking Δ*t* → 0 gives the continuous-time version:

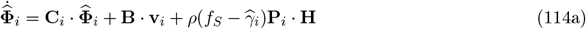

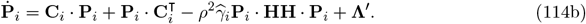

Here, the discrete-time feedback *f*_*S,t*_ has converged to a sum of delta functions *f*_*S*_, and

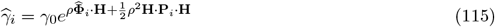

is the estimated feedback rate.

equations (114a) and (114b) correspond to equations (58a) and (60a) in the continuous feedback case, with *ρ***P**_*i*_ · **H** playing the role of feedback gain **K**_*i*_, and 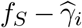 playing the role of the prediction error. Whereas the optimal gain scales with the precision of feedback observations in the continuous case, here it scales with the steepness of the feedback rate function.

As above, the feedback gain can be approximated by partitioning the covariance matrix **P**_*i*_ (see equation (66)), and ignoring the minor contribution of *x*_*i*_ to the lower-right block,

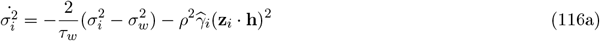

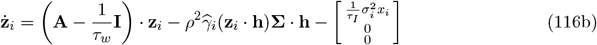

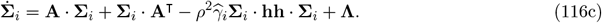

A key difference between these equations and equation (69) is that in the latter **Σ**_*i*_ relaxes to a steady state, whereas here it depends dynamically on the estimate of the feedback rate 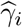. Intuitively, the higher the feedback rate, the more uncertainty about the error is reduced.

The rest of the derivation proceeds identically to the continuous case. Unpacking equation (114a) into the original notation, and setting 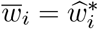, yields an error estimate, 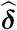, covariance, **Σ**, and feedback rate estimate, 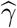, that are all independent of the synapse index. The control gain is computed identically to that above (equations (61) and (65)), yielding a control variable 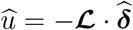.

Putting everything together, the plasticity rule for spiking feedback is

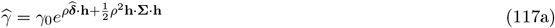

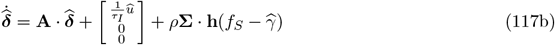

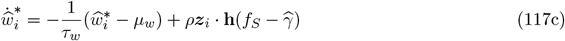

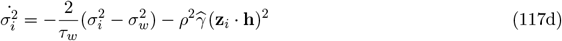

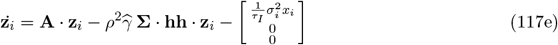

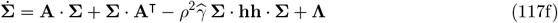

with weights set via equation (77) as before.

### Feedback delays

In the brain, error feedback cannot be provided instantaneously to a neuron due to communication and sensory processing delays, typically on the order of ∼ 10 − 100 ms. We model this constraint by introducing a lag of time *τ* into the feedback signal, modifying equation (10) to become

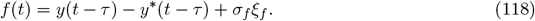

Similarly, for the spiking feedback model, equation (17) becomes

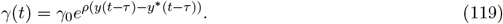

For the slow weights, the learning rules can be adapted by simply feeding lagged copies of the input and weight parameters into the governing equations, such that the Bayesian filter estimate reflects the unobserved states as they were at time *t* − *τ*. Working through the details, the only change required to the learning rules is to replace *x*_*i*_(*t*) with *x*_*i*_(*t* − *τ*) in equation (76d), and similarly in the spiking feedback case. Such a lagged copy of the input could be maintained at the synapse via simple chemical signaling cascades, as shown by Jayabal et al. (2022). As long as the assumed target-weight drift is slow relative to the feedback delay, *τ*_*w*_ ≫ *τ*, this simple modification of the learning rule maintains consistent performance in the presence of delays.

For the fast weights, we employ the Smith predictor technique to build a controller that accounts for the delay (Smith 1959). Conceptually, we need to compute a control signal at time *t* that will cancel the current error, but we only have an estimate from feedback observations of the error as it was at time *t* − *τ*. We therefore use the known model dynamics and recent history of applied control signals to predict the current error by propagating the estimate ***δ*** forward in time. We use this to construct the controller

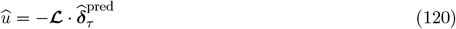

and thus the fast-weight rule

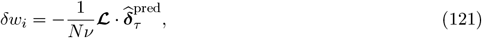

where the control gain ℒ is the same as that above.

Writing 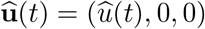, the predicted error can be computed via

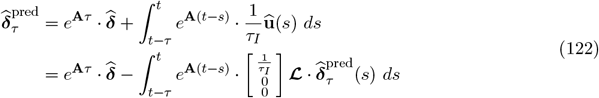

The first term propagates 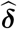 forward in time via a matrix exponential that encodes the deterministic dynamics of the error vector. The second term is a convolution that accounts for the influence of past controls – propagating the corrections applied by the controller at each time *s* (whose effects have not yet been observed) to the current time *t*.

The convolution can be simplified by considering an online implementation. Denoting the integral by **U**(*t*), differentiation gives

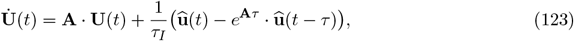

which only requires knowing the most recently applied control and a single lagged variable from the past.

### Multiple neurons

In Fig. 3C and Fig. 4, we simulated a small feedforward network of *M* neurons. In this model, each of the neurons has identical dynamics, as described by equations (7a) and (7b), but receives independent synaptic input and noise. Noise variances were scaled by a factor 1*/M* relative to the single neuron simulations, for ease of comparison (see Table 1 for parameters). A common downstream output is driven by the sum over firing rates, *r*_*m*_, described by equation (19). A spiking feedback signal, *f*_S,*m*_, is generated independently for each neuron using the Poisson rate function in equation (17).

In the plasticity rule, the current, rate and synaptic noise terms in equation (71) are scaled by *M* to reflect the contribution of all neurons to the error. The fast weight rule (equation (46)) also needs to be modified to account for the fact that the feedback signals, and therefore error estimates, differ across neurons. Ignoring locality constraints, the optimal strategy would be to combine the error estimates computed by each of the *M* neurons, 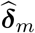, and their uncertainties, **Σ**_*m*_, as a precision-weighted average,

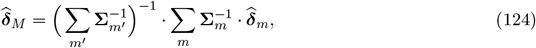

and use that to construct a single global control variable, 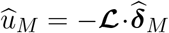. We use a local approximation of this strategy, by replacing the precision weighting in equation (124) with a scaled identity matrix,

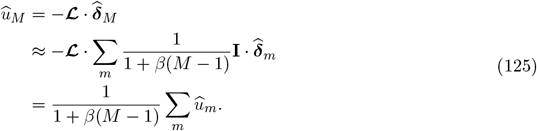

In equation (125), 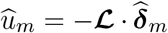 is the control variable computed locally by neuron *m*, and *β* ∈ [0, 1] is a free parameter (fitted in pilot simulations to minimize task output error). When feedback rates are very low, a small number of neurons would dominate the sum in equation (124) at any given time (error estimates are most precise immediately following a feedback spike, but relax to small, uncertain baseline values soon afterwards), so *β* should be small. In the limit of very high feedback rates, all terms would be weighted equally, so *β* ≈ 1. Intuitively, *β* interpolates between sparse and dense feedback regimes. Using this approximation leads to scaled fast weights

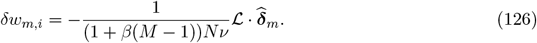

### Simulation details

#### Online linear regression

For the simulations in Fig. 1, the model neuron was driven with sinusoidal inputs and required to learn weights that would produce a time-varying, periodic target output. We used *N* = 20 synapses and a period *T*_period_ = 1 s.

Input rates were defined as

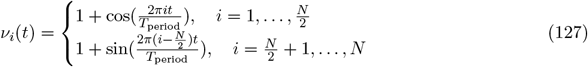

The target output was generated as

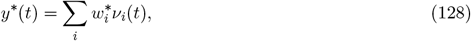

With 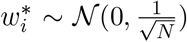. Learning rules were as described by equations (2), (4) and (6), with learning rates 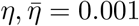.

### Teacher-student task

In Figs. 2 and 3, simulations were initialized by independently drawing target weights 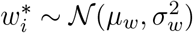, input rates *ν*_*i*_ ∼ uniform(0, *ν*_max_), slow weights 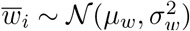, and setting fast weights *δw*_*i*_ = 0. The weight uncertainties 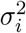 were initialized with value 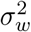. Input spikes were generated at each time step by drawing binary variables *x*_*i*_ ∼ Bernoulli(*ν*_*i*_Δ*t*). Feedback spikes were generated similarly in the spiking-feedback case, using the rate function in equation (17), and also enforcing a hard maximum of 200 spikes*/*s for stability. Target weights drifted slowly about the mean *µ*_*w*_, as described by the Euler discretization of equation (41). Actual weights were updated according to the Euler discretization of equation (76) in the continuous case, equation (117) in the spiking-feedback case, and analogously for the classical rules. Performance was quantified by computing the root-mean-squared error between output and target output over all time points in the simulation, thus measuring both learning speed and steady-state error.

### Cerebellar learning task

The cerebellar learning task in Fig. 4 requires mapping time-varying patterns of synaptic input to associated target outputs. Unlike the teacher-student task, the target outputs are generated as random wavelets, rather than through explicit target weights.

Input patterns of duration *T*_pattern_ = 1 s were constructed in a similar manner to Bicknell and Häusser (2021). A synapse was selected to be active on a given pattern with probability *p*_active_ = 0.5. If active, its time-dependent presynaptic firing rate in that pattern was defined by drawing a peak time *t*_peak,*i*_ ∼ Uniform(0, *T*_pattern_), maximum amplitude *ν*_max,*i*_ ∼ Uniform(90, 110) (spikes/s) and centering a Gaussian bump of activity within the pattern interval,

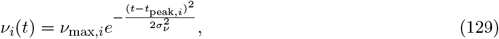

with width parameter *σ*_*ν*_ = 0.1 s. Input spikes were generated stochastically each pattern presentation using this rate function.

For the target outputs, we generated random trajectories that drift slowly over time and could be stitched together in random order to yield a single continuous target (analogous to composing different sequences of motor actions). This was implemented using truncated Fourier series with random coefficients that drift via Ornstein-Uhlenbeck processes. Different trajectories are forced to agree at the endpoints by using a common constant offset and multiplying the time-dependent parts by a bump function. Specifically, the drifting target output for pattern *p* was defined via

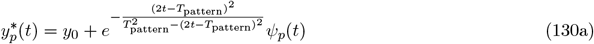

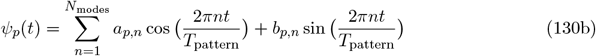

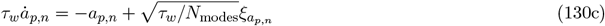

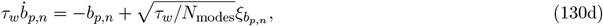

with initial coefficients *a*_*p,n*_, 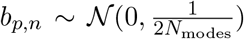. We used parameters *N*_modes_ = 3, which sets the maximum temporal frequency, and *τ*_*w*_ = 10^4^ s, equal to the weight drift parameter in the spiking-feedback teacher-student task.

### Plasticity protocol

For the simulated plasticity experiments in Fig. 5, we used models that had been trained on the cerebellar learning task. We applied the protocol to 100 synapses using the classical and Bayesian learning rules, initializing with the trained weights and uncertainties. The classical learning rate, *η*, and control cost, *λ*_*u*_, were set at the values that minimized output error during the task. For most simulations, the protocol consisted of repeated pairings of parallel fiber input bursts (5 spikes at 10 ms intervals) and single climbing fiber spikes at a given delay. The numbers of spikes were varied in a subset of simulations (Fig. S3c,d). The pairing was repeated 50 times at 500 ms intervals. Plasticity effects were computed at the end of the protocol as the change in slow-weights (or classical weights) from their initial trained values.

## Code availability

Simulation code is available at https://github.com/babicknell/SynControl

## Acknowledgments

This work was supported by the Gatsby Charitable Foundation and the Wellcome Trust (110114/Z/15/Z). The authors acknowledge the use of the UCL Myriad High Performance Computing Facility (Myriad@UCL), and associated support services, in the completion of this work.

## Supplementary Figures

**Figure S1:**
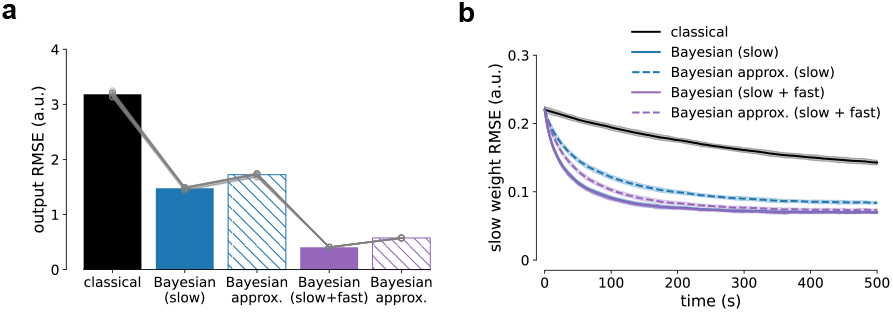
Approximation of the Bayesian learning rule. Comparison of the full Bayesian rule, given by equation (76), and the approximation presented for ease of interpretation in equations (13) and (16), in terms of output error (**a**) and weight error (**b)**. The approximation is valid whenever the output time constant dominates the model dynamics, *τ*_*r*_, *τ*_*I*_ ≪ *τ*_*y*_. Here we use parameters *τ*_*I*_ = 1 ms, *τ*_*r*_ = 10 ms, and *τ*_*y*_ = 100 ms.

**Figure S2:**
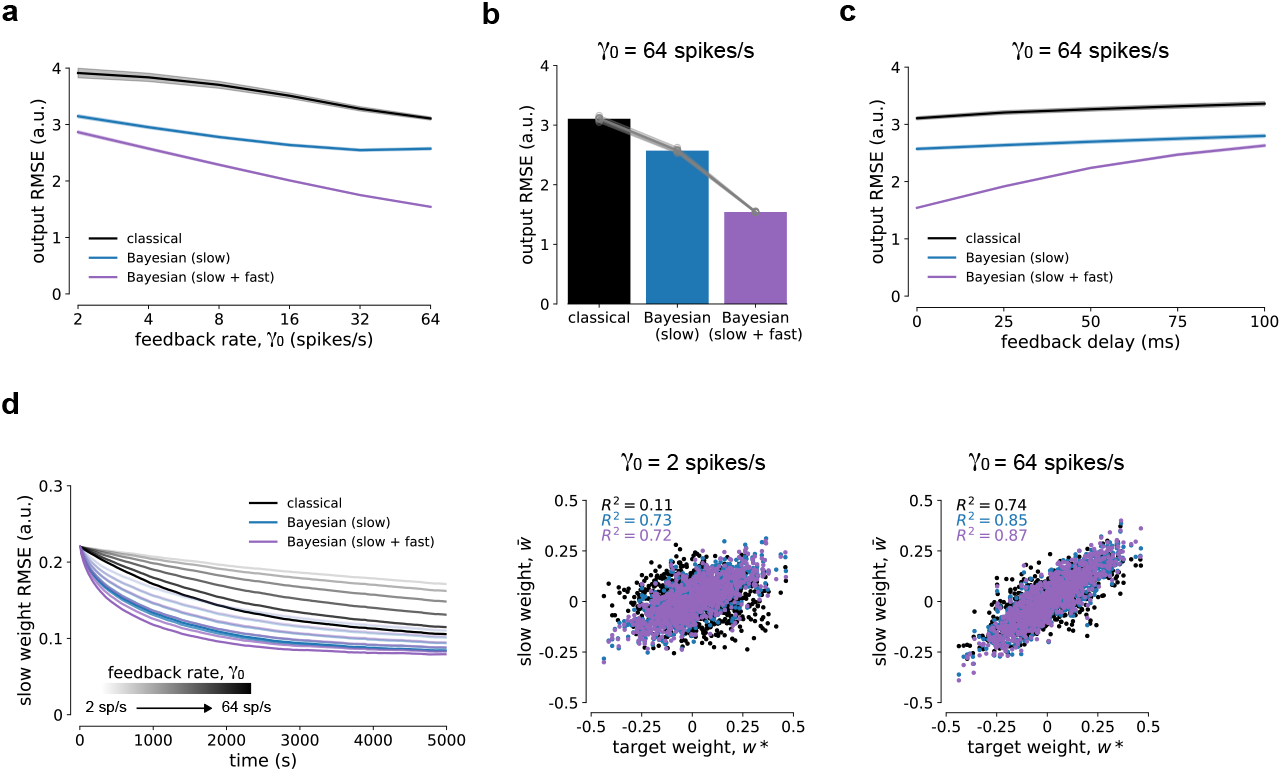
Simulations with spiking feedback. Results from the spiking feedback model are consistent with the continuous feedback model (Fig. 2), although learning takes longer and performance depends strongly on the spontaneous feedback rate *γ*_0_. **a)** Performance as a function of feedback rate (replotted from Fig. 3 for context). Shaded areas are s.d. from 10 random seeds. **b)** Detailed comparison between learning rules with *γ*_0_ = 64 spikes/s. **c)** Control remains effective in the presence of feedback delays. Shaded areas are s.d. from 10 random seeds. **d)** Weights converge to their target values more quickly with increasing rates of feedback.

**Figure S3:**
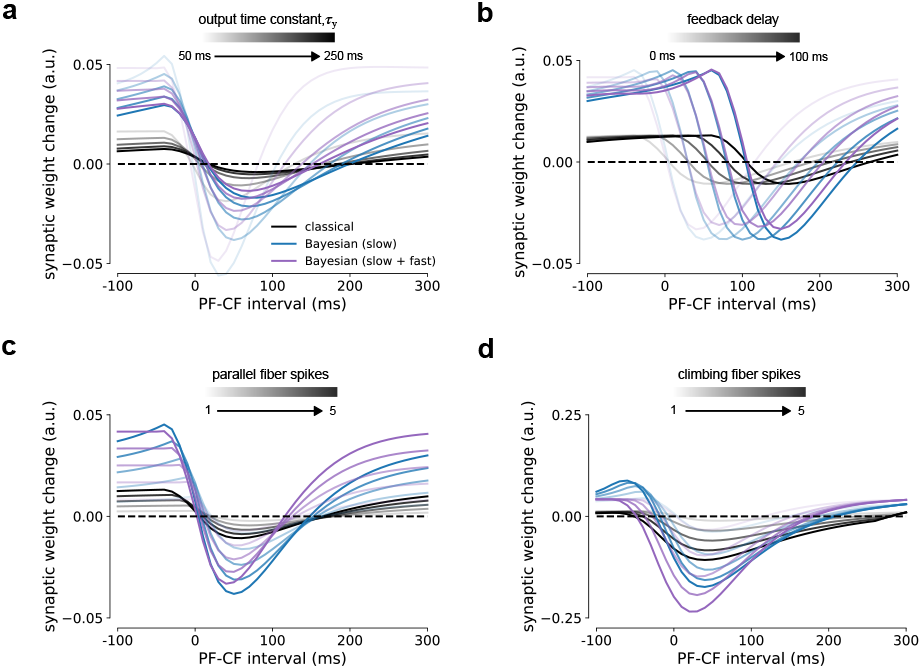
Parameter dependence of simulated plasticity experiments. The LTP/LTD curve produced using the protocol in Fig. 5c depends systematically on the parameters of the model and protocol. **a)** Tuning to the PF-CF interval becomes broader as the output time constant *τ*_*y*_ increases. **b)** Peaks in the LTP and LTD lobes of the curve are shifted in proportion to the feedback delay. **c)** The magnitude of plasticity increases with the number of spikes in the parallel fiber burst. **d)** LTD is amplified and shifted to earlier PF-CF intervals with increasing numbers of climbing fiber spikes.

